# A lipid nanoparticle platform for high yield CRISPR-targeted homology directed repair enables fully non-viral CAR T cell generation

**DOI:** 10.64898/2026.08.04.741347

**Authors:** Joy J Chen, Atip Lawanprasert, Alissa LW Kleinhenz, Mehar Gayatri Devi Namala, Daniel Chu, York Tang, Valentine Lindarto, Michael Launspach, Danna Lee, Hoa Wu, Niren Murthy, David N Nguyen

## Abstract

CRISPR-mediated homology directed repair (HDR) enables targeted CAR integration with improved fitness and therapeutic potential of CAR T cells. However, current methods for generating HDR-engineered CAR T cells rely on viral transduction or electroporation, approaches that limit global implementation and constrain patient access due to their cost, toxicity, and requirement for centralized manufacturing. Through a screen of ionizable lipids, we identified LNP systems that enable CRISPR-mediated gene knock-in (KI) in primary human T cells and are amenable to hand mixing by ethanol injection as a research tool or machine formulation for larger scale manufacturing. Modifying the linear dsDNA HDR template with truncated Cas9 target sequences (tCTS) enhanced HDR rates across multiple LNP systems. We optimized two LNP formulations capable of HDR-mediated KI of a large 4kB CD19 CAR-EGFR HDR template into the TRAC locus with rates of ≥8% and >10x improved edited cell yields compared to electroporation. We demonstrate that LNP-generated CAR T cells exhibited similar growth kinetics, activation states, differentiation states, and killing capacity compared to electroporation-generated CAR T cells. Our LNP platform components are fully disclosed, commercially sourced, and enable efficient fully non-viral CRISPR-HDR cell engineering across diverse applications.

## Main Text

The first CAR T cell products were approved by the FDA in 2017, and to date seven CD19-and BCMA-targeting CAR T cell products have been approved for various hematologic malignancies. However, despite this progress, a substantial gap remains between the number of CAR T-eligible patients and those who receive CAR T therapy due to the prohibitive cost of good manufacturing practice (GMP)-grade viral vectors, complex logistics, and manufacturing constraints associated with CAR T cell production^1,2^. All seven FDA-approved CAR T products rely on centralized manufacturing using integrating lentiviral or retroviral vectors, and the resulting 3-to 6-week turnaround time further contributes to patient mortality^1,3^. This access gap is even more pronounced globally, as low-resource settings are particularly vulnerable to the financial toxicity of current CAR T cell manufacturing pipelines^1,4,5^.

The semi-random integration of CAR transgenes caused by lentiviral or retroviral vectors carries risk of insertional mutagenesis and creates substantial variability in CAR expression ^6–8^. CRISPR-mediated targeted integration of a CAR transgene into the T cell receptor alpha (*TRAC*) locus endogenously regulates CAR expression with improved function and reduced T cell exhaustion^8^. However, the delivery methods required for homology directed repair (HDR) in primary T cells presents manufacturing challenges^3^. The standard protocols rely on EP^8,9^ to deliver Cas9 ribonucleoproteins (RNPs) editors and DNA repair templates. However, the high-voltage pulses required for efficient DNA delivery induce substantial cellular stress, resulting in acute cytotoxicity, delayed proliferation, and impaired T cell phenotype^10,11^. To achieve comparable cell yields to those obtained with viral manufacturing methods, EP requires higher starting cell numbers and longer expansion times. This poses as a challenge for patients when coupled with low apheresis yields in hematologic cancers and the critical need to minimize vein-to-vein time^12^. While adeno-associated viruses (AAVs) for secondary HDR delivery can be coupled with CRISPR editor delivery by EP^8,13^ or other non-viral strategies^11^, AAVs are costly at the required manufacturing scale and the prevalence of anti-AAV antibodies limits future *in vivo* potential^10,14,15^.

Self-assembling lipid nanoparticles (LNPs) offer a promising strategy to overcome these manufacturing and toxicity bottlenecks, serving as an attractive cell manufacturing alternative to both viral and EP-based engineering strategies^16^. While LNPs can efficiently deliver RNA cargoes in most cell types^17^, efficient nuclear delivery of the long DNA templates required for HDR has been elusive outside of tumor cell lines^18,19^. Further, LNP-delivered CRISPR-mediated HDR in primary human T cells has previously required a hybrid strategy relying on AAV delivery of donor templates^10^.

Here, we present a fully non-viral, all-LNP platform for high-efficiency, CRISPR-targeted CAR knock-in (KI) into the *TRAC* locus of activated primary human T cells using commercially available readily sourced reagents. Starting with a screen of ionizable lipids, we developed LNP formulations that achieved HDR in T cells with efficiency and functionality matching EP while avoiding its cellular toxicity, resulting in substantially higher viable T cell yields. In addition, these LNPs facilitated delivery of a wide range of payloads, from small ssODNs (0.1kb) to large linear dsDNA HDR templates (1.4-4kb) and circular single-stranded DNA (cssDNA) formats (1.3-2.2kb)^20^. The efficacy of dsDNA was enhanced in multiple LNPs by incorporating truncated Cas9 target sequences (tCTS)^21,22^. Coupling the ease of formulating LNPs with superior edited cell yields compared to electroporation, the versatility of LNPs supports the potential of delivery of a variety of payloads tuned to specific cell therapy applications.

## Results

### LNPs based on the ionizable lipid L-319 enable CRISPR-HDR in T cells

Although many proprietary LNPs can deliver mRNA to primary T cells, it remains unclear whether they can efficiently co-deliver Cas9 mRNA, guide RNA, and HDR template to enable targeted KI. To develop an LNP-only platform that enables high-yield CRISPR-targeted HDR in primary human T cells, we first screened a library of LNPs made with different ionizable lipids. We assembled a chemically diverse library of ∼70 commercially available ionizable lipids with pKa values between 6.0 and 7.0, which balances LNP stability with potency of endosomal disruption^18,23^. LNPs were manually formulated by ethanol injection^24^ and screened in batches across 4 unique blood cell donors for delivery of GFP mRNA to activated T cells (**Supplementary Fig. 1-2**). We identified 12 unique LNP formulations, which transfected T cells with GFP mRNA with high efficiency and reasonable viability. These hits were further screened for delivery of CRISPR reagents in activated primary human T cells. LNPs were co-formulated by ethanol injection with a *RAB11A* sgRNA, Cas9 mRNA, and a 1.4kb linear dsDNA HDR template encoding KI of an N-terminal GFP-RAB11A fusion (RAB11A-GFP HDR template) **(Fig. 1a)**. The sgRNA, Cas9 mRNA, and HDRT DNA were added at a 1:1:1 mass ratio and mixed at a 10:1 total lipid: nucleic acid mass ratio. Flow cytometry was used to assess GFP expression indicating successful KI at day 3. The L5 LNP, based on the biodegradable L-319 ionizable lipid^25^, emerged as a top candidate from our library screen achieving ∼5% RAB11A-GFP^+^ KI with low toxicity **(Fig. 1b-d).** For further experiments, we also included LNPX, a recently published clinically relevant lipid for mRNA delivery^26^ based on the SM-102 ionizable lipid, which also demonstrated high *B2M* disruption and RAB11A-GFP^+^ KI yields (**Supplementary Fig. 3**).

**Figure 1.**
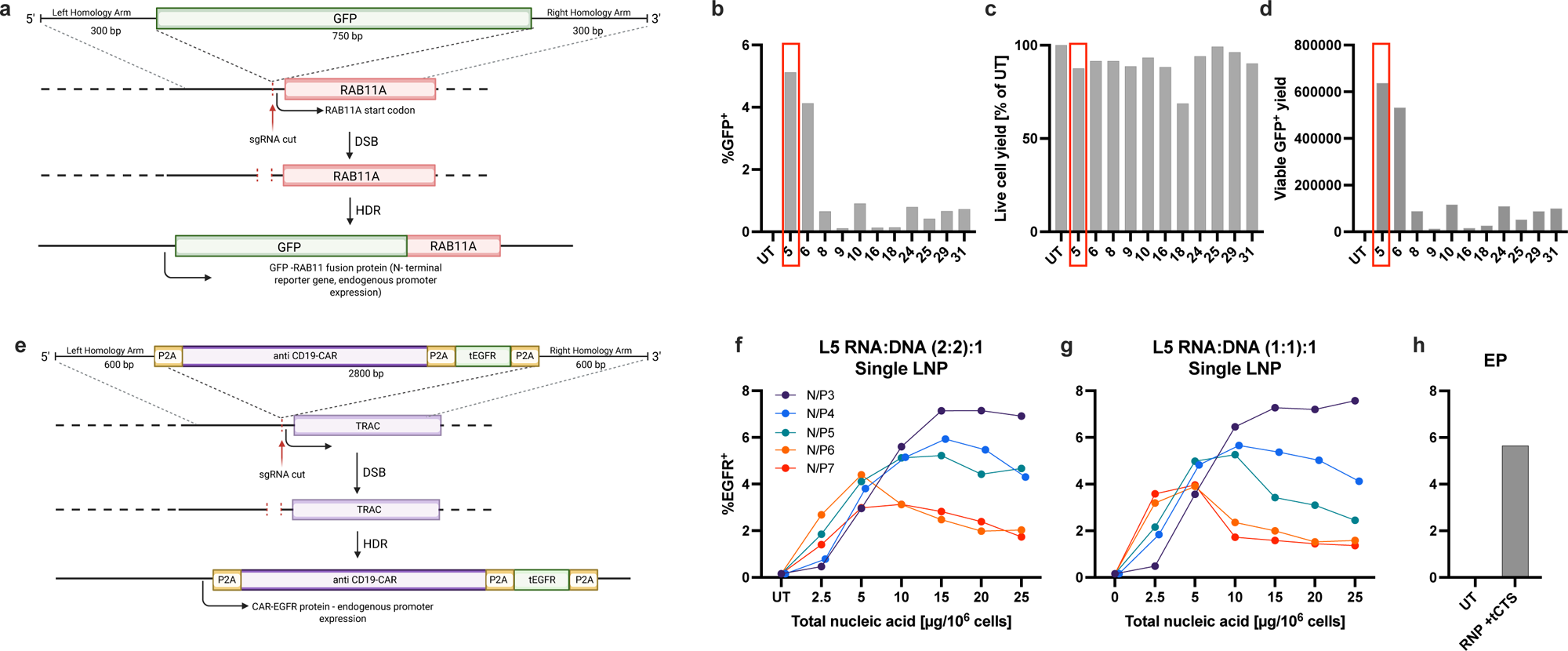
LNPs based on the ionizable lipid L-319 generate HDR in primary T cells with an efficiency comparable to electroporation. (a-d) Ionizable lipids identified from the high-throughput GFP screen (Supplementary Fig. 2) were assessed for HDR-based gene editing in activated primary human T cells at the *RAB11A* locus. **(a)** Schematic of the ∼1.4 kb dsDNA template encoding sfGFP flanked by *RAB11A* specific LHA and RHA LNPs were formulated by ethanol injection to co-encapsulate *RAB11A* sgRNA, Cas9 mRNA, and RAB11A-GFP HDR template and were delivered to primary activated human T cells. Day 3 flow cytometry was used to assess **(b)** editing efficiency, shown as the percentage of GFP^+^ cells, **(c)** live cell yield, expressed as a percentage of untreated controls, and **(d)** edited GFP^+^ viable cell yield. *N* = 1 biological donor. **(e-f)** Candidate lipid L5 was advanced to assess KI of large dsDNA CAR-EGFR payload. **(e)** Schematic of the ∼4 kb donor DNA template encoding anti-CD19 CAR and tEGFR reporter separated by a P2A self-cleaving peptide and flanked by *TRAC*-specific LHA and RHA. Day 3 flow cytometry was used to assess CAR-EGFR editing efficiency of ethanol injection LNPs at varied varying N/P and RNA:DNA ratios co-encapsulating Cas9 mRNA, *TRAC* sgRNA, and CAR-EGFR HDR template at (sgRNA:Cas9 mRNA):DNA w/w/w ratios of **(f)** (2:2):1 or **(g)** (1:1):1. **(h)** *TRAC* RNP and CAR-EGFR HDR template with tCTS delivered via EP. UT, untreated; tCTS, truncated Cas9 target sequences; LHA, left homology arm; RHA, right homology arm; *N* = 1 biological donor.

We hypothesized that LNP delivery of large HDR constructs may alleviate the compounding toxicity effect from both electroporation and the HDR template. We optimized the L5 LNP formulation for delivery of a clinically relevant 4kb HDR template encoding *TRAC*-targeted anti-CD19 CAR with co-expressed truncated EGFR (TRAC-EGFR HDR template) **(Fig. 1e)**. While HDR rates are typically inversely proportional to the size of the insertion construct, toxicity seems directly proportional to HDR template length^12,27^. An initial optimization of LNP formulated with ethanol injection was performed by combinatorial screening across key parameters (N/P ratio, RNA:DNA ratios, and total nucleic acid dose) in a single donor. The N/P ratio, defined as the molar ratio of ionizable lipid amines (N) to nucleic acid phosphates (P), is a critical formulation parameter known to influence particle formation, encapsulation efficiency, and delivery efficacy^28^. We observed this to be true in LNPs formed by ethanol injection and found that for L5, the highest HDR rates occur at low N/P ratios and at RNA:DNA ratios with lower DNA content. While we observed similar HDR rates between EP and LNP in the same donor, we found lower toxicity in the LNP condition, consistently yielding higher CAR-T cell numbers compared to EP when normalized to the input number of cells (**Fig. 1f-h, Supplementary Fig. 4**). Although we observed performance variation across both LNP batch and cell donor, these results highlight primary human T cell CRISPR HDR delivery that can be achieved with hand-mixed LNPs without special equipment.

### LNPs formulated by microfluidic mixing can generate CAR T cells that express a large CD19 CAR-EGFR via CRISPR-HDR

We further optimized parameters for automated microfluidic mixing of LNPs, which can improve particle quality, batch-to-batch consistency, and delivery potency compared to ethanol injection^29^. Using the commercially available NanoAssemblr™ Spark™ formulation system, we tested L5 and LNPX LNPs for encapsulation of both RNA and DNA across various N/P ratios and nucleic acid doses. We observed narrow particle size polydispersity by DLS (**Supplementary Fig. 5a-c**) and high encapsulation efficiencies determined via Ribogreen (**Supplementary Fig. 5d-f**).

We independently investigated delivery of either RNA or DNA containing LNPs by encapsulating and testing each component separately. RNA-only LNPs (Cas9 mRNA + sgRNA) targeting three different genes (*B2M*, *CD5*, and *TRAC*) exhibited comparable editing efficiency compared to optimized EP conditions in freshly PBMC-isolated T cells **(Supplementary Fig. 6a-c)**; previously frozen T cells had similar optimal RNA doses but with slightly lower editing efficiencies **(Supplementary Fig 6d-f).** While LNPX is more potent than L5, at optimal dosing of either L5 or LNPX we observed similar yields of LNP-edited knock-out cells compared to EP **(Supplementary Fig. 6a-f).** As cytosolic dsDNA causes cellular toxicity^30^, we attempted to minimize the total HDR template dose required to facilitate HDR at the *RAB11A* locus with the RAB11A-GFP HDR template **(Fig. 1a)**. We approached this with a dual LNP KI approach, varying the ratio of RNA:DNA while keeping the total dose constant to evaluate the impact of each component. We observed expected high KI rates with optimal dosing for EP^21^, while LNP-treated T cells had lower rates. Increasing the ratio of DNA:RNA, while maintaining the same total nucleic acid dose, does not significantly impact the overall KI percent or yield (**Supplementary Fig. 7).**

Having established parameters for efficient KI at *RAB11A* locus, we further optimized LNP composition and culture media conditions for delivery of a larger therapeutically relevant 4kb CD19 CAR–EGFR dsDNA construct^8^ **(Fig. 1e).** We varied L5 formulations comparing the helper lipids (DOPE and DSPC), cholesterol analogues (β-sitosterol)^31^, and media conditions (+/-fetal bovine serum (FBS) and +/-Apolipoprotein E (ApoE)). In contrast to previously reported LNPX formulations for mRNA delivery using β-sitosterol and no ApoE at time of treatment^26^, L5 transfection was most optimal using DOPE helper lipid, standard cholesterol (**Supplementary Fig. 8)**, serum-free culture at time of LNP addition, and required ApoE for optimal transfection (**Supplementary Fig. 9)** indicating low-density lipoprotein receptor (LDLR)-mediated uptake^32^. With these established parameters, we next confirmed optimal dosing for CD19-CAR HDR, observing greater potency with LNPX than L5 but both achieving higher yield of successfully edited cells than standard RNP-HDR template KI by EP **(Fig 2a-d).** Performing a similar optimization across RNA:DNA ratios like for the RAB11A-GFP KI, we observe that increasing the relative amount of encapsulated DNA for the larger CAR-EGFR HDR template does not improve overall KI or edited cell yield, but all conditions resulted in KI percentages equivalent to or higher than EP **(Supplementary Fig. 10).**

**Figure 2.**
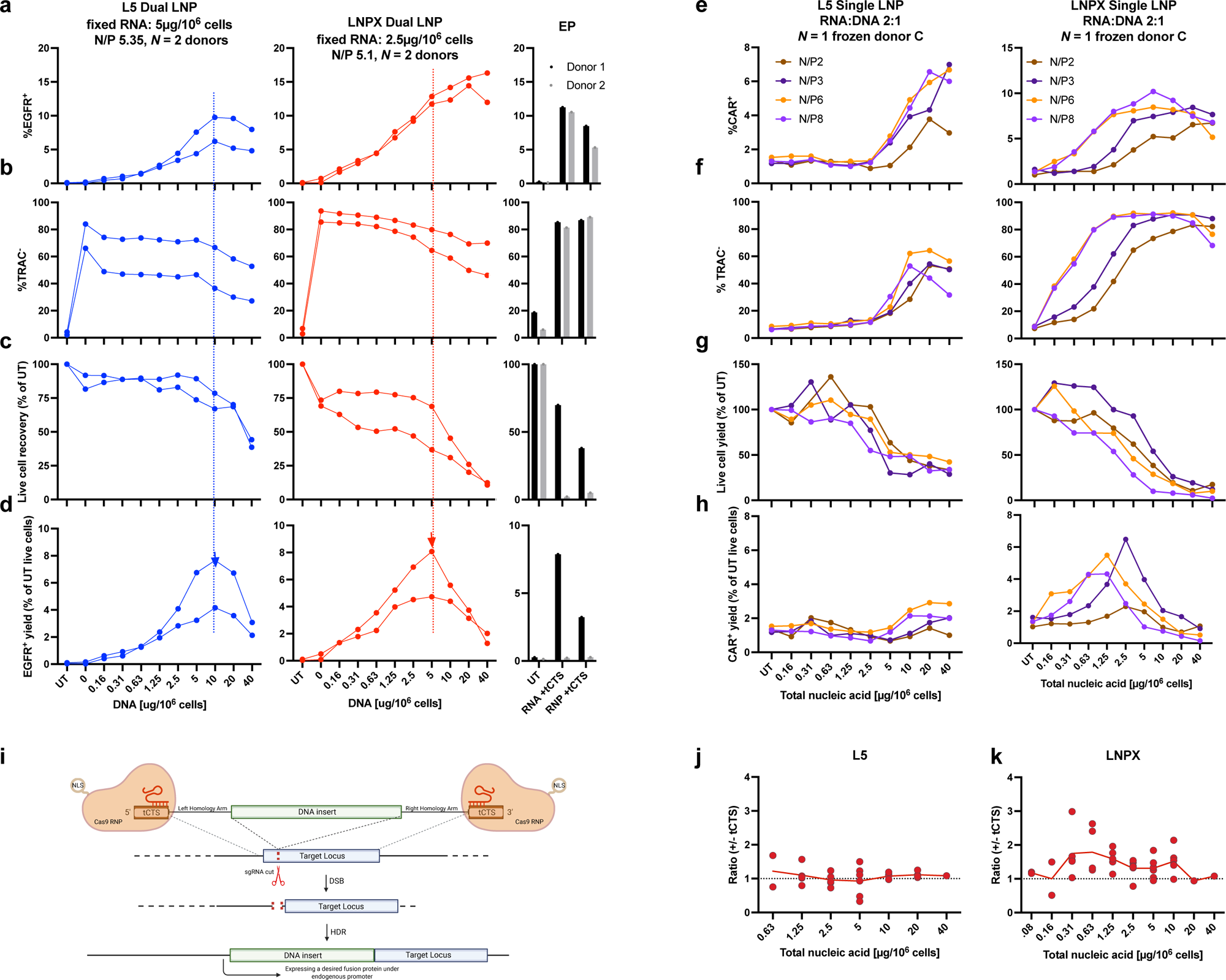
Microfluidic LNP formulation optimization highlights optimal LNP dose for high editing efficiency and tCTS-mediated HDR. LNPs containing *TRAC* sgRNA, Cas9 mRNA, and a CAR-EGFR HDR template with tCTS were formulated either by **(a-d)** dual encapsulation of RNA and HDR in separate LNPs (*N=2)* or **(e-h)** single encapsulation of RNA and DNA components (*N=3)*. LNP parameters were optimized by **(a-d)** DNA dose with constant optimal RNA dose and by **(e-h)** NP ratio with a constant 1:1:1 w/w/w sgRNA:Cas9mRNA:HDR template ratio. Day 3 flow cytometry was used to assess **(a, e)** KI efficiency, shown as the percentage of EGFR^+^ or CAR^+^ cells, respectively. (b, f) *TRAC* disruption, shown as the percentage of *TRAC*^-^ cells. **(c, g)** Live cell yield, expressed as a percentage of untreated controls. **(d, h)** Edited cell yield, calculated as the product of relative live cell recovery (normalized to UT) and EGFR^+^ or CAR^+^ frequency, respectively. **(i)** Schematic of tCTS mechanism. tCTS allows HDR template to bind to RNP that have NLS that can bring the HDR template into the nucleus. **(j-k)** CAR-EGFR KI was compared in L5 and LNPX LNPs containing HDR templates with or without tCTS over a range of doses with sgRNA:Cas9mRNA:HDR template ratio w/w/w of 1:1:1. Data summarize single-encapsulation and dual-encapsulation experiments (*N* = 1-6). UT, untreated; tCTS, truncated Cas9 target sequences; NLS, nuclear localization signal.

### A single particle all-in-one LNP generates CAR T cells that express a CD19-CAR with similar efficacy to CAR T cells generated from dual encapsulation

Although independent LNP encapsulation of RNA and DNA cargoes allows granular control of encapsulated components at the bench, single encapsulation facilitates manufacturing and future *in vivo* CAR-T engineering. Comparing single vs dual encapsulation for our payloads at selected doses, we found edited CAR-T cell yields to be comparable across the various encapsulation strategies **(Supplementary Fig. 11).** We further screened HDR efficiency for all-in-one LNPs mixed in a NanoAssemblr™ Spark™ across a range of N/P ratios, surprisingly revealing that unlike manual-formulated LNPs by ethanol injection, a higher N/P ratio of 6-8 demonstrated superior performance across *N* = 3 unique cell donors (**Fig 2e-h**, **Supplementary Fig. 12)**. This difference may partially be attributed to reduced excess lipid in the uniform machine-mixed LNPs as ethanol injection leaves excess lipid species that could induce toxicity at higher N/P ratios.

### Optimizing the DNA payload improves LNP-mediated HDR efficiency

Cytoplasmic delivery is sufficient for mRNA applications, but nuclear entry of DNA is a critical barrier for achieving CRISPR-targeted HDR by LNP delivery following endosomal escape^18,19^. We incorporated truncated Cas9 target sequences (tCTS) into the linear dsDNA HDR templates as a basis for screening and optimization of LNPs, based upon their proven success in electroporation^21,22^. These tCTSs facilitate the interaction between the HDR template and the Cas9 RNP which allows the Cas9 nuclear localization signal (NLS) to promote HDR^21^. We included two tCTS sequences on all HDR templates as a baseline and we tested if removing tCTSs from the HDR templates had an impact upon LNP-mediated HDR rates. We observed the benefit of keeping tCTSs across multiple LNP platforms, particularly lower-efficiency manually formulated LNPs made by ethanol injection (**Supplementary Fig. 13).** In more optimized machine formulated LNPs, there was no detriment to including the tCTS sequences in KI efficiency for short ssODN encoding KI of an HA tag at the *CD5* locus^22^ (**Supplementary Fig. 14)**. In most donors, the tCTS boosts HDR at several RNA:DNA ratios for both L5 and LNPX in the 1.4kb RAB11A-GFP HDR template (**Supplementary Fig. 7, 15)** and in the 4kb CAR-EGFR HDR template (**Supplementary Fig. 10, 16 and Fig 2j-k)**. Using dual particle LNP systems we also tested if timing of the DNA-LNP addition to cells affected HDR outcomes, and we observed tCTS-containing HDR templates to be most beneficial at add simultaneously with the RNA-LNP, consistent with early nuclear delivery facilitating HDR at the time of initial DSB generation **(Supplementary Fig. 17).**

We hypothesized that alternative DNA formats and shorter HDR template lengths may provide benefits for mitigating toxicity. Circular single stranded DNA (cssDNA) has emerged as low-cytotoxicity alternative DNA format for non-viral gene therapy^33^ including delivery of HDR templates by EP in primary cells *ex vivo*. We tested a ∼1.3kb RAB11A-GFP cssDNA template in L5 and LNPX compared to linear dsDNA with or without tCTS (**Supplementary Fig. 18)**. Both LNPs can encapsulate cssDNA and have comparative KI and yield compared to linear dsDNA (**Supplementary Fig. 18)**. We also tested a ∼2.2kb CD19 CAR cssDNA template (no EGFR transgene, 300bp HAs) in LNPX using the same formulation parameters optimized for linear dsDNA and found HDR rates comparable to delivery by EP with higher overall yield by EP (**Supplementary Fig. 19**). We further tested LNP delivery of HDR templates with shorter homology arms (450bp compared to 600bp) on the linear dsDNA CAR-EGFR HDR template and observed that both dsDNA conditions with 600bp or 450bp homology arms performed similarly with higher edited cell yield when compared to EP **(Supplementary Fig. 19).**

### LNP-mediated CAR T cell generation improves cell yield across culture conditions and supports scale-up

Across multiple donors and experiments, LNP editing generated orders-of-magnitude greater live cell yield and CAR T-cell yield per input cell than EP when evaluated under modality-specific conditions: low density culture for LNP delivery (10k cells/ 200µL in a 96-well plate) and high-density culture for EP (200k cells/200µL in a 96-well plate well) **(Fig. 3a-b)**. LNPs generated marked improvement in overall live cell and CAR^+^ cell yields in the previously frozen donors, reflecting greater EP-associated toxicity and consequently lower EP yields. Pooling donors and experiments across single-versus dual-encapsulation revealed comparable high CAR^+^ T cell yields, supporting scale-up manufacturing (**Supplementary Fig. 20)**. To directly compared EP-and LNP-generated CAR T cell yields at matched culture conditions and higher input cell numbers to assessed scalability for manufacturing, we tested three batch sizes: small (200µL in 96 well plates), medium (2mL in 12 well plates), and large (40mL in T-75 flask) **(Fig. 3c)**. While editing rates were similar across conditions in these donors (∼5% by EP and 5-8% by LNP), both L5 and LNPX all-in-one formulations exhibited less cytotoxicity and consistently outperform EP for total edited CAR+ T cell yields in all tested culture densities and batch sizes **(Supplementary Fig. 21, Fig. 3c)**. Delivery with LNPX generated 60x more CAR-T cells in the 96-well format and 13X more CAR-T cells in the 12-well format compared to EP, suggesting these LNP platforms are robust for scaling batch *ex vivo* manufacture of primary human T cells toward cell therapy applications **(Fig. 3a-c, Supplementary Fig. 20).**

**Figure 3.**
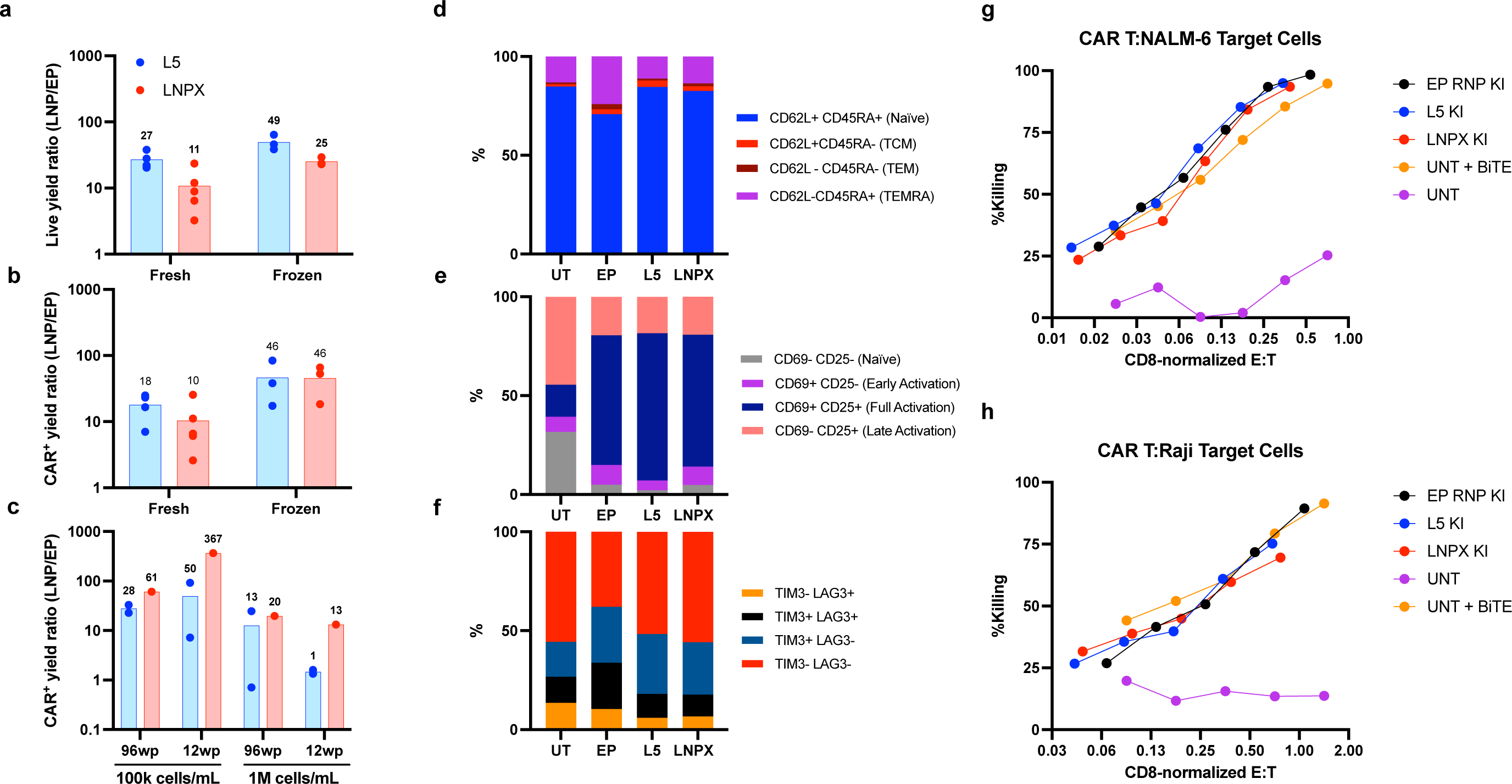
LNP-engineered CAR T cells supports scale-up, preserve edited CAR T phenotype compared to electroporation, and exhibit robust cell killing activity. (a-c) Comparison of LNP/EP ratio of live cell yields and editing yields assessed by Day 3 flow cytometry. **(a)** Live cell yield ratio for L5-and LNPX-edited cells. **(b)** CAR^+^ cell yield ratio for L5-and LNPX-edited cells. Ratios were calculated as the yield per input cell following LNP delivery divided by that following EP. Data are from *N* = 4-5 fresh donors and *N* = 3 previously frozen donors, cultured at optimal conditions for each editing method. **(c)** CAR^+^ yield cell ratio for L5-and LNPX-edited cells cultured at matched culture conditions across each LNP and EP. **(d-f)** T cell phenotyping assessed prior to cell killing assay using flow cytometry on Day 17 post edit showing **(d)** memory states with CD62L and CD45RA, **(e)** activation states with CD69 and CD25, and **(f)** exhaustion with LAG3 and TIM3 flow markers. **(g-h)** L5 and LNPX generated CAR-T cells have indistinguishable killing capacity in a luminescent cell killing assay compared to EP CAR T and UT + bispecific T cell engager (BiTE) in both **(g)** CD19+ NALM-6 cells and **(h)** CD19+ Raji cells. *N* = 1 biological donor. UT, untreated; tCTS, truncated Cas9 target sequences.

### LNP-engineered T cells preserve edited CAR T cell phenotypes and function

Scaling up the established LNP editing systems in a 12-well plate at the optimal dose for dual encapsulation for L5 (5ug RNA/10^6^ cells and 10ug DNA/10^6^ cells) and LNPX (2.5ug RNA/10^6^ cells and 5ug DNA/10^6^ cells), we generated CAR-T cells and evaluated their functional and phenotypic profiles. EGFR-based magnetic enrichment was used to purify EGFR^+^ CAR T cells from ∼3% to >90% (**Supplementary Fig. 22)**. We compared the growth kinetics of LNP-generated and EP-generated CAR T cells and observed a faster growth rate in the LNP-edited samples than EP-edited samples pre-enrichment, but similar growth profiles after EGFR enrichment at day 10 of expansion **(Supplementary Fig. 23).** T cell subsets were evaluated via markers of T cell memory (CD62L and CD45RA), activation (CD69 and CD25), and exhaustion (TIM3, LAG3) by flow cytometry on day 17 post edit. We observed that LNP-generated CAR T cells maintain similar memory subsets and activation states as EP-generated CAR Ts and untreated T cells **(Fig. 3d-f).** Finally, enriched CAR-Ts were evaluated for their cytotoxic potential in a targeted cell killing assay by co-culturing them with either CD19+ NALM-6 or CD19+ Raji cell lines (**Supplementary Fig. 24)**. Normalizing for subtle shifts in the CD8 fraction **(Supplementary Fig. 25**), we observed no functional differences in cytotoxicity between L5 and LNPX LNP-generated and EP-generated CAR T cells, achieving near-complete target cell lysis at high CD8 effector:target (E:T) ratios after 20 hours of co-culture (**Figure 3g-h).** After the cell killing assay, the exhaustion profile was also similar across LNP-and EP-generated CAR-Ts, with a slight increase in CD25 activation and slight decrease in CD45RA+CD62L+ naïve cells post CD19 co-culture (**Supplementary Fig. 26).** Taken together, these findings suggest that while both delivery methods impose some degree of cellular stress, LNP-mediated engineering achieves greater edited cell yields while maintaining similar T cell proliferation and functional outcomes compared to the EP standard.

## Discussion

Despite the transformative efficacy of engineered T cell therapies, widespread clinical use is constrained by complex manufacturing processes that we begin to address with a fully non-viral LNP-based system for CRISPR editing in primary human T cells. We optimized two LNP formulations, L5 based upon the L-319 ionizable lipid^25^ and LNPX based upon the SM-102 ionizable lipid^34^. Both LNPs enable CRISPR-targeted HDR KI in primary human T cells of templates ranging in size from 0.1-4 kb, including a complex CD19 CAR construct with a co-expressed EGFR safety switch. Remarkably, despite screening 70 commercially available ionizable lipids, the two LNPs that emerged for T cell engineering are two of the most highly engineered and well-studied ionizable lipids even though they were developed for distinct purposes: L-319 for small RNA delivery^25^ and SM-102 for large mRNA delivery^35^. Comparatively, we find that LNPX achieves higher KI rates than L5 at the tradeoff of higher toxicity. L-319 has an ester group instead of the parent molecule DLin-MC3-DMA’s (MC3) 9,10-cis double bond, which introduces a hydrolytically labile linkage cleaved under physiologic conditions and may result in relatively less stability than SM-102^25^. L-319 also demonstrates rapid clearance in animal models within a few hours, with an estimated half-life of ∼30 minutes in serum^25^. Consequently, L5 may be better suited for lab-scale and ethanol injection applications where preparation-to-treatment timeframes are shorter and lower overall toxicity may be prioritized. In contrast, LNPX may be preferable where higher % editing is required or for GMP-scale manufacturing. They also each have different mechanisms of cell uptake with L5 requiring ApoE utilizing the low-density lipoprotein receptor (LDLR)^32^, while LNPX transfection is ApoE-independent^26^.

The accessibility of LNP systems contrasts with the conventional *ex vivo* HDR T cell engineering pipelines. These workflows rely on either (i) EP of Cas9 RNPs with dsDNA templates, which requires specialized electroporation equipment and training^21,36^ prohibitive for decentralized or resource-limited settings or (ii) hybrid AAV6 systems to deliver DNA templates^10^, which is restricted by a ∼4kb packaging limit and high GMP manufacturing burdens^37^. Instead, we demonstrate that our LNP systems can be readily formulated at the bench without specialized instrumentation enabling standard research labs to use this method of gene delivery for many applications. Continued development with microfluidic mixing systems and GMP-scale implementation of all-LNP HDR platforms could substantially expand access to CRISPR-engineered cellular therapies, as LNPs represent a cost-effective and accessible alternative to the existing clinical *ex vivo* cellular therapy manufacturing workflows dependent on expensive and genotoxic recombinant viral vectors. Further, LNP transfection is operationally similar to viral transduction, meaning current GMP-compliant cell therapy manufacturing infrastructure (including closed-loop systems for local manufacture^3^)could be adaptable to LNPs^36^.

As development of LNP-HDR systems progress, several limitations should be acknowledged. As expected, we see donor-to-donor variation in HDR rates and toxicities. Efficiency of HDR is dependent upon cell cycle as optimal expression of homologous recombination machinery occurs during S and G2 phases^38,39^. All studies were performed in activated healthy donor T cells, which is a typical limitation of CRISPR-HDR editing in primary T cells; even with optimal activation, we observe HDR efficiency remained donor-dependent. Development of T cell targeting leading to activation is likely to be required to ultimately facilitate resting T cell HDR editing that will enable *in vivo* applications. A tradeoff is expected between DNA dose enabling CRISPR-HDR and toxicity^21, 27^. Intriguingly, HDR rates are preserved with LNPs even with lower relative DNA doses, suggesting that total donor DNA dose is not a constraining variable.

Instead, editing may reflect the sum of DNA’s opposing actions depending on its cellular localization: cytoplasmic DNA activates the innate immune system via the cGAS-STING pathway and contributes to toxicity^30^, while nuclear DNA is the actual HDR substrate. Some studies show cGAS-STING signaling can inhibit HDR^30^, which could support why LNP delivery achieves HDR even at low doses while at higher doses increased cytoplasmic DNA eventually decreases HDR rates. To address the inherent limitations in nuclear delivery of DNA by LNPs, we added tCTS-enhanced HDR templates achieving modest but reproducible benefit across multiple LNP platforms (**Fig 2j-k, Supplementary Fig. 14-17**) but with wide variation across biological donors. Surprisingly, the tCTS HDRTs did not impart additional toxicity in the LNPs, which had previously been observed for tCTS HDRTs delivered by electroporation^21,22^. Thus, while tCTS HDRTs do not always help in the context of saturating doses of LNPs *ex vivo* (particularly L5), they were rarely detrimental and may be best suited for improving HDR rates at low LNP-DNA doses whereby any individual cell has minimal transfection events – a delivery regime likely to be encountered *in vivo*.

A central finding of this study is that LNP-mediated delivery approached HDR frequencies of optimized electroporation workflows without the major viability penalties typically associated with electroporation. Both LNPX and L5 generated ∼10-50x higher numbers of CAR T cells **(Fig. 3a-b)** particularly in previously-frozen cells utilized in most clinical workflows, with more rapid expansion in the first week of culture **(Supplementary Fig. 23)** while maintaining phenotypic and functional characteristics comparable to EP-generated CAR T cells **(Fig 3d-f)**. Thus, beyond permissive manufacturing, LNPs may have additional additional functional benefits for rapid manufacturing of CAR-Ts, permitting a rapid manufacturing approach that minimizes *ex vivo* culture times with clinical benefit^3,40^, and may be a better approach to achieving a therapeutic dose of CAR-T cells particular for patients recently exposed to lymphodepleting chemotherapies. Future studies will be required to evaluate performance in quiescent cells, patient-derived material, and GMP-scale manufacturing workflows.

For LNP optimization experiments, T cells in LNP conditions were cultured at lower initial densities (10k-50k cells per 200uL) than electroporated cells (133k-200k per 200uL). LNP-mediated transfection depends on maximizing nanoparticle-cell interactions for cellular uptake^41^ so are often cultured at lower concentrations, whereas electroporation results in substantial acute cell loss^27^ and is followed by culture at higher cell concentrations to support recovery and expansion. When comparing EP and LNP-generated CAR T generation across different cell culture densities and scales, LNPs consistently yielded ≥10 fold greater numbers of successfully edited CAR T cells **(Fig 3b)**. Though the improvement in cell yields was more modest (1.3x for L5, 13x for LNPX) at larger culture formats; KI frequencies remained high at all scales **(Supplementary Fig. 21).** Although conditions were chosen to match across volumes or surface areas, additional differences may be due to cell distribution and medium depth in the different vessel geometries and require future vessel-specific optimization. The CAR-EGFR HDR template used is approximately 4kb and further work is required to determine the upper limits of payload for LNP-mediated HDR delivery. While we see KI efficiencies at levels comparable to optimized electroporation systems, HDR efficiency and clinical scale manufacturing robustness could be further improved via template optimization.

As a vehicle for nucleic acid medicines, LNPs established a favorable clinical safety profile having been administered in billions of vaccines does and other therapeutic applications^42^ without the insertional mutagenesis risks of viral vectors or emergence of anti-vector immune responses^43^. Beyond simplifying manufacturing and the prospects for expanding cell therapy access, our all-in-one non-viral LNP platform establishes design principles that may ultimately support HDR-mediated *in vivo* CAR-T generation. This concept is distinct from current *in vivo* approaches that either use LNPs to deliver CAR mRNA, resulting in transient receptor expression, or use engineered lentiviral vectors to install a CAR transgene through semi-random, non-locus-targeted integration. In contrast, CRISPR-catalyzed HDR-mediated CAR insertion into the *TRAC* locus yields regulated and durable receptor expression during T cell proliferation while reducing effector differentiation and exhaustion^8^. Although significant delivery challenges remain before clinical application of HDR-based *in vivo* CAR engineering becomes feasible, our findings support the possibility that fully non-viral systems may eventually enable such approaches.

In summary, we have developed fully non-viral LNP systems optimized for CRISPR-targeted HDR in primary human T cells at high efficiencies as a generalizable framework for delivery of DNA payloads including CAR-T generation. A modular platform in which a common LNP formulation can be paired with different guide RNAs and HDR templates could accelerate development of diverse cellular therapies while maintaining a shared manufacturing workflow. Importantly, we disclose the commercially available ionizable lipids used in our formulations (L5 with L-319 and LNPX with SM-102) and the LNP compositions we optimized. This provides an open framework for others to adapt quickly and build upon for diverse applications. In principle, such an approach would not only support CAR T cell products, but also gene-corrected autologous T cell therapies for a broad range of monogenic immune disorders and other engineered cell therapy applications.

## Online Methods

### Cell culture

Primary adult blood cells from anonymous healthy male human donors were purchased as leukapheresis packs (STEMCELL #200-0092), and bulk T cells were isolated by magnetic negative selection using EasySep isolation kit for CD3+ T cells (STEMCELL, #19051) per manufacturer protocol. T cells were either used fresh or stored in liquid nitrogen until use. Isolated CD3+ bulk T cells were cultured in X-VIVO 15 (Lonza) supplemented with 5% FBS and 50 uM 2-mercaptoethanol (ie X-VIVO+). Fresh cells were activated on day of isolation (day-2 or day-3 relative to treatment) and cultured at 1e6 cells/mL in X-VIVO+ with anti-human CD3/CD28 magnetic DynaBeads (Gibco, #40203D) at a bead to cell ratio of 1:1 and a cytokine cocktail of 300 U/mL IL-b (ThermoFisher, AF-200-02-1MG), 5 ng/mL IL-7 (ThermoFisher, AF-200-07-100UG), and 5 ng/mL IL-15 (ThermoFisher, AF-200-15-100UG); or 10ng/ml IL-7 and 10ng/mL IL-15 but no IL-2. Previously frozen cells were first thawed in non-supplemented X-VIVO 15 for 4-24h, then activated in the manner described for 72h (day-3 relative to treatment). On day of treatment, Dynabeads were removed from cell culture by diluting 1:1 v/v with buffer (PBS without calcium/magnesium supplemented with 10% FBS and 1 mM EDTA), vigorous pipetting, then isolating cell suspension after 6 minutes incubation on a EasySep cell separation magnet (STEMCELL, #18002) to remove magnetic beads.

### sgRNA Preparation

Alt-R™ sgRNA (IDT) were resuspended in Nuclease Free Duplex Buffer (IDT, #11-01-03-01) to a concentration of 80µM for making RNP. They were then further diluted with Nuclease Free Duplex Buffer to 1µg/µL for LNP formulation. sgRNA stocks were stored in-80C once resuspended and used for up to 5 freeze-thaw cycles. The TRAC sgRNA used in the yield comparison experiments (Fig 3a, Supplementary Fig. 21) had Intellia modifications.

### Ribonucleoprotein (RNP) preparation

To make RNPs, a volume ratio of 1:0.8:1 of 80 µM sgRNA, 100 mg/mL polyglutamic acid (PGA), and 40uM Cas9 protein was prepared and incubated at 37C for 15 min, as previously described [**31819258**]. Once formed, RNPs were frozen and stored in-80C and used after no more than 1 freeze-thaw cycle. Unless otherwise stated, the final dose of RNP per nucleofection was 50 pmol on a Cas9 protein basis.

### HDRT template preparation

RAB11A-GFP HDRT and CAR-EGFR HDRT plasmids were amplified via PCR with tCTS containing primers, as previously described^21^designed for the cognate guide RNA used for KI editing HDRT. Guide and primer sequences listed in **Supplementary Table 2 (Excel)** and HDRT sequences listed in **Supplementary Table 3 (Excel)**. Gel electrophoresis was performed to confirm the template size and no evidence of high molecular bands or smearing indicative of concatemer formation. The HDRT amplicon was purified using 0.8X AMPure XP Beads (Beckman Coulter, #A63882) per manufacturing recommendations and eluted in nuclease-free water. Spectrophotometry (A260) was performed by NanoDrop (Thermo Scientific, #13-400-527) to determine the concentration of HDRT. Short ssODN HDRT templates were synthesized as Ultramer oligonucleotides (IDT) and resuspended to 100uM. To anneal tCTS sequences, the ssODN was mixed with tCTS oligos at a 4 oligo:1 ssODN ratio. The mix was heated up to 95C and cooled by 1C per minute using a thermocycler, as previously described^22^. A circular single stranded DNA (cssDNA) HDR template encoding the CD19-CAR insertion targeted into the *TRAC* locus was cloned into a proprietary vector backbone, produced, and isolated by Full Circles Therapeutics^20^, with cssDNA concentration verified by absorbance (Nanodrop). Cloning, Sanger, and Nanopore sequence verification was performed by Quintara Biosciences. CssDNA encapsulation in LNPs was performed with the same formulation parameters and NA ratios as for linear dsDNA HDR templates. All HDRTs were stored in-20C before use.

### T cell electroporation

Electroporation was performed on the Lonza 4D-Nucleofector System in a 96 well plate format. Frozen RNPs were thawed and incubated at 37C for 5 minutes. HDR templates were mixed and incubated with RNPs for at least 5 min prior to addition to 96 well nucleofection plates (Lonza). Unless otherwise stated, 50pmol RNP, 1-1.25 µg Cas9 mRNA, 1-1.25 µg sgRNA, and/or 0.5-1ug HDR template were used per million cells. Immediately prior to electroporation, T cells were centrifuged for 9-10 min at 100 x g, media was aspirated via vacuum, and cells were resuspended in electroporation buffer P3 (Lonza) using 16–20 μl buffer per 0.5–1.0e6 cells. 0.5-1e6 cells were pipetted into the 96 well nucleofection plate and gently mixed with RNP +/-HDRT. Cells were electroporated with pulse code EH-115 for RNP +/-HDRT or with pulse code DS-137 for mRNA-sgRNA editor. Cells were rescued in 80µL X-VIVO+ (Lonza, 02-060Q) supplemented with 10% FBS immediately post electroporation and incubated for 10 minutes in 37C before seeding into XVIVO+ supplemented with 100-250 U/mL IL2 and 10% FBS and cultured at a concentration of 0.5-1E6 T cells/mL.

### EGFR positive enrichment

Edited cell enrichment was performed using the EasySep™ Human EGFR Positive Selection Kit, following manufacturer protocol (STEMCELL, #100-1131).

### LNP treatment

T cells were seeded at 10,000-50,000 cells per well in 50 µL in a 96-well plate in treatment media (serum-free X-VIVO supplemented with 250-500U IL2 and 1µg/mL ApoE3). LNP treatments were diluted to the appropriate concentration in 50 µL treatment media and gently mixed into cell culture. Cell cultures were topped off with 100 µL X-VIVO+ after 24-48 hours. Experiments done in 12 well plates were seeded at 500k cells in 500 µL and topped off with 500 µL after 24-48 hours. LNP treatments were spiked into the 500 µL volume, instead of pre-diluted in media.

### Flow cytometry

Cells were collected for flow cytometry on day 3 or 4 post-transfection and washed with MACs buffer (1% FBS and 1µM EDTA in PBS) and centrifuged at 300 x g for 5 minutes before cell surface staining with 30µL antibody mix and incubating for 15-30 minutes on ice in the dark before washing. Staining panel included viability dye of GhostDye 780 (Cytek, #13-0865-T500) or Fixable Near IR Viability Dye (Fisher Scientific, #L34982), AF488 EGFR (Biolegend, #352908), PE TCRa/b (Biolegend, #306708), or PE G4S Linker (Cell Signaling Tech, #38907S) and APC TCRa/b (Biolegend, #306718). Full antibody list described in **Supplementary Table 1.** Flow cytometry was performed on the Attune NxT or CytoFLEX LX Flow Cytometer. All flow cytometry data was analyzed using FlowJo software v10.10.0 (BD Biosciences).

### Cell lines

NALM-6-GFP-Luciferase and Raji-Luciferase cell lines were provided by the laboratory of Julia Carnevale at the University of California, San Francisco (UCSF). Both cell lines were cultured in RPMI 1640 medium (ThermoFisher Scientific, #11875093) supplemented with 10% FBS and 1% penicillin-streptomycin.

### Cell killing assay

EGFR-enriched effector CAR T cells were seeded from 20,000 cells per well with serial dilutions to achieve a final range from 2:1 to 0.0625:1 E:T ratio; 10,000 CD19+ target cells (either NALM-6 or Raji) were added to each well. CAR T cells and target cells were diluted in and cultured in 100 µL RPMI 1640 (ThermoFisher Scientific, #11835030) with no phenol red and no IL2 for 20 hours. An anti-CD19-anti-CD3 bispecific molecule was added to unedited cells as a positive control for cell killing (BPS Biosciences, #100441-1). Luminescence was quantified after adding LucScreen (Thermofisher Scientific, #T1033) on a Tecan Spark High performance Multi-Mode Microplate Reader, controlled via SparkControl software (Tecan).

### Manual LNP formulation by ethanol injection for GFP mRNA LNP screening

Curated LNP formulation candidates were screened for their transfection efficiency in activated primary human T cells by delivering GFP mRNA. Screened LNPs were generated by ethanol injection, the rapid mixing of the organic phase containing each lipid mixture into the aqueous phase containing GFP mRNA (0.33 mg/mL in 25 mM acetate buffer, pH = 4). A fixed ratio of 3:1 aq.-to-organic and 10:1 lipid-to-nucleic acid weight ratios was used. Each Lipid mixture was generated by combining a unique ionizable lipid with helper lipid DOPE (Avanti Polar Lipids), cholesterol (ChemScene), and DMG-PEG 2000 (Avanti Polar Lipids) at a fixed volumetric ratio of 4:3:2.5:0.5. A complete list of ionizable lipid candidates used to create lipid mixtures is provided in **Supplementary Table 4 (Excel).** All lipids were prepared in ethanol at a concentration of 10 mg/mL. After ethanol injection, LNPs were incubated at room temperature for 10 minutes before transferring the LNP-mRNA mix with an equivalent of 100ng mRNA into 10,000 cells/well of primary human T cell culture in 100µL XVIVO+ supplemented with 1µg/mL ApoE3. 24 hours after treatment, cells were stained with a viability dye (GhostDye 780) then analyzed on an Attune Nxt Flow cytometer for GFP expression.

### Manual LNP formulation by ethanol injection for CRSIPR-Cas9 mediated HDR LNPs

LNPs encapsulating RAB11A, B2M, CD5, or TRAC, Cas9 mRNA (Trilink), and respective HDR template were encapsulated in the LNP with the previously described methods above, with the mass ratio of 1:1 sgRNA:Cas9 mRNA (for gene knock-out) or 1:1:1 sgRNA:Cas9 mRNA: DNA (for gene KI). Cells were treated with 125 ng total nucleic acid per 10,000 T cells **(Supplementary Fig. 3)** or at the listed RNA:DNA ratio and doses, indicated in the figures and legends. At indicated timepoint post-transfection, cells were washed then stained for flow cytometry to quantify editing efficiency.

### Automated microfluidic LNP manufacturing

LNPs used for CAR-T production were manufactured by microfluidic mixing (NanoAssemblr Spark, Cytiva). Nucleic acid cargoes were prepared in a 25mM acetate buffer, pH = 4, to a final nucleic acid concentration of 0.25 mg/mL. L5 organic phase was prepared by mixing 50% L-319, 10% DOPE, 38.5% Cholesterol, and 1.5% DMG-PEG2000 in ethanol to the final concentration of 10 mg/mL. LNPX organic phase was prepared by mixing 50% SM-102, 10% DSPC, 38.5% β-sitosterol, and 1.5% DMG-PEG2000 in ethanol to the final concentration of 10 mg/mL. LNPs were formulated by the NanoAssemblr^TM^ Spark^TM^ microfluidic system or Ignite^TM^ following manufacturer’s protocol at a 20:1 nucleic acid:lipid ratio, unless otherwise stated and immediately diluted into 1X DPBS. The crude LNPs were purified by diafiltration using an Amicon-0.5 filter with a molecular weight cutoff of 100 kDa into DPBS buffer. Final DNA and RNA concentrations were quantified using Quant-iT Ribogreen RNA Kit (ThermoFisher Scientific, #R11490) which is a fluorescent dye-based assay to quantify nucleic acid. Standards were prepared from the aqueous phase of the LNP mix. LNP were plated in duplicate under two conditions: 1. Untreated to measure free, unencapsulated RNA or DNA, and 2. treated with 2% Triton X-100 to disrupt LNPs and release encapsulated cargo. Encapsulated cargo concentration was calculated as the difference between total cargo measured after Triton X-100 disruption and free cargo measured in untreated samples. Encapsulation efficiency was calculated as [(total cargo (Triton X-100) – free cargo) / total cargo (Triton X-100)] * 100. LNPs were then added to pre-activated T cells as described above with final dosing as indicated. Dynamic Light Scattering (DLS) was performed on a Malvern DLS Zetasizer to track LNP size across batches for quality control.

## Plots and Schematics

Figure plots were made in GraphPad Prism.

Schematics and illustrations were created with BioRender.Com.

## Acknowledgements

The CD19-CAR-EGFR construct was a gift from the lab of Prof. Justin Eyquem at the University of California San Francisco.

## Author Contributions

J.J.C, A.L., A.L.W.K., N.M. and D.N.N. conceived of the study. J.J.C, A.L., A.L.W.K., M.G.D.N., D.C., Y.T., N.M., D.N.N. designed experiments. J.J.C, A.L., A.L.W.K., M.G.D.N., Y.T., D.C., V.L., contributed to the completion of experiments. J.J.C, A.L., A.L.W.K., M.G.D.N., and M.L. analyzed the data. D.L. and H.W. provided the cssDNA from FullCircles Therapeutics. J.J.C., A.L.W.K., A.L.L., M.G.D.N., N.M., and D.N.N. wrote the manuscript. All authors read and approved the final manuscript.

## Funding acknowledgements

This research was funded, in part, by a gift from the Medical Excellence Research Innovation Trust (MERIT) philanthropic fund. This research was funded, in part, by the Advanced Research Projects Agency for Health (ARPA-H), an agency within the U.S. Department of Health and Human Services (HHS), under Award Number 140D042590005. The views and conclusions contained in this document are those of the authors and should not be interpreted as representing the official policies, either expressed or implied, of the U.S. Government. D.N.N. was supported by NIH grants L40AI140341, K08AI153767. J.C. was supported by the National Science Foundation Graduate Research Fellowship under Grant No. DGE 2146752. Any opinion, findings, and conclusions or recommendations expressed in this material are those of the authors(s) and do not necessarily reflect the views of the National Science Foundation. A.L.W.K was supported by the National Institute of Diabetes and Digestive and Kidney Diseases of the National Institutes of Health under Award Number TL1DK139565. The content is solely the responsibility of the authors and does not necessarily represent the official views of the National Institutes of Health. M.L. was supported by the Deutsche Forschungsgemeinschaft (DFG, German Research Foundation) through the Walter Benjamin Programme (Project-ID 571675766).

## Availability of data and materials

All data produced in the present study are available from the corresponding author upon reasonable request.

