## Supplementary Figures and Table 1 for "A lipid nanoparticle platform for high yield CRISPR-targeted homology directed repair enables fully non-viral CAR T cell generation"

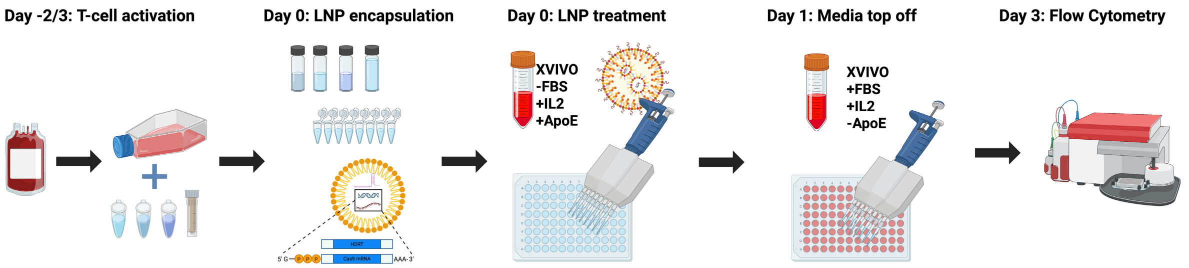


**Supplementary Figure 1. Schematic of LNP- based gene delivery in activated primary human T cells.** Bulk T cells were isolated from healthy donors and activated for 2–3 days prior to LNP treatment using anti-human CD3/CD28 magnetic Dynabeads™ at a bead-to-cell ratio of 1:1, supplemented with a cytokine cocktail of either 300 U/mL IL-2, 5 ng/mL IL-7, and 5 ng/mL IL-15, or 10 ng/mL IL-7 and 10 ng/mL IL-15. LNPs were formulated either via ethanol injection or by using the NanoAssemblr™ Spark™ or Ignite™ on the day of treatment. Cells were seeded at a density of 10k - 50k cells/well in 50 µL of serum-free X-VIVO 15 supplemented with either 250 - 500 U/mL IL-2 and 1 µg/mL ApoE, or 10 ng/mL IL-7, 10 ng/mL IL-15, and 1 µg/mL ApoE. LNPs were diluted to the appropriate concentration in 50 µL treatment media and added to cells. After 24 - 48 hours post-treatment, cells were topped up with 100 µL X-VIVO 15. On Day 3, cells were assessed for both editing efficiency using flow cytometry.

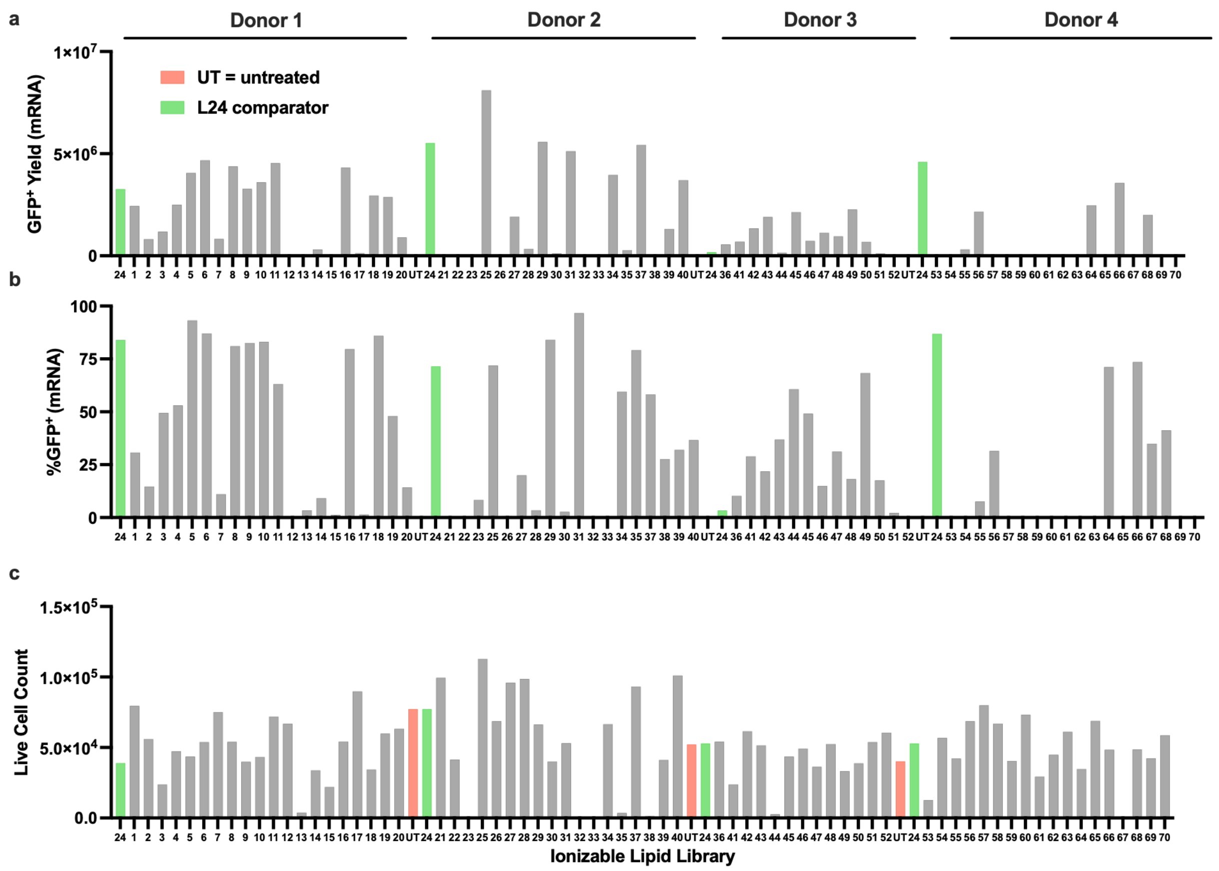


**Supplementary Figure 2. Screen of 70 ionizable lipids for GFP mRNA delivery.** The assembled lipid library was evaluated in four biological donors using L24 as an internal comparator due to its consistently high baseline GFP expression. LNPs were formulated by ethanol injection encapsulating GFP mRNA. Day 2 flow cytometry was used to assess **(a)** GFP$^+$ cell yield, **(b)** % mRNA transfection (%GFP$^+$ cells), and **(c)** total live cell count. UT, untreated; *N* = 1 biological donor.

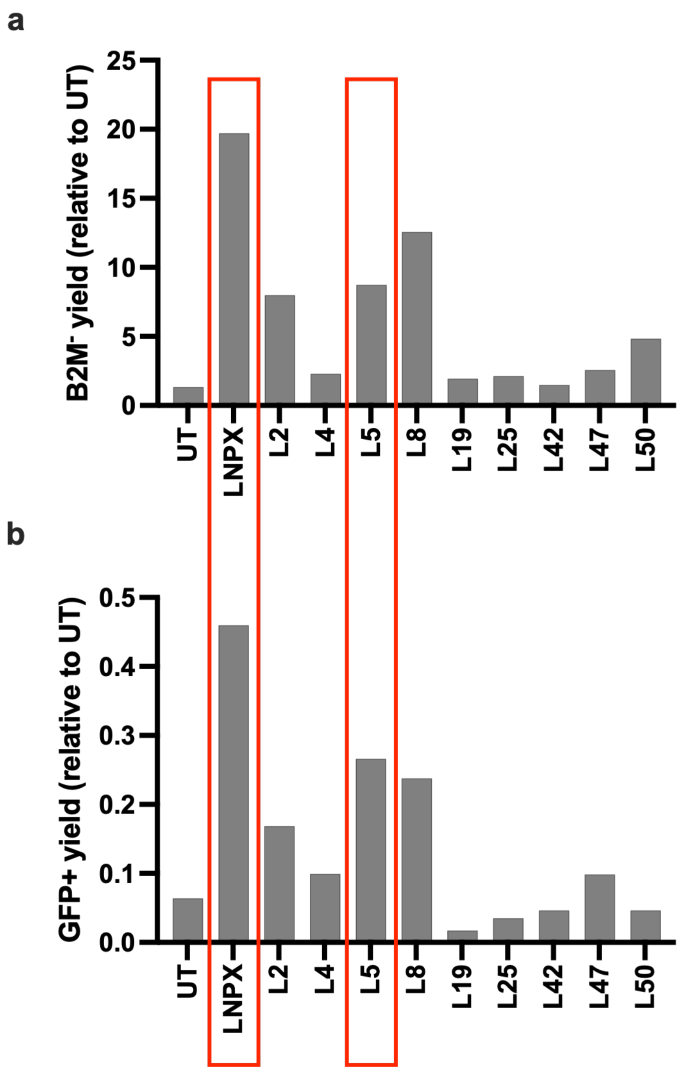


**Supplementary Figure 3: Screening LNP candidates for delivery of CRISPR editing machinery.** LNPs co-encapsulating either B2M sgRNA and Cas9 mRNA or RAB11A sgRNA, Cas9 mRNA, and RAB11A-GFP HDR template +tCTS were formulated by ethanol injection. Day 3 flow cytometry was used to assess cell counts representing cell yield relative to UT of **(a)** gene disruption at B2M and **(b)** RAB11A-GFP knock in. Yield was calculated as the product of live cell recovery (normalized to UT) and edited cell frequency. UT, untreated; tCTS, truncated Cas9 target sequence, *N* = 1 biological donor.

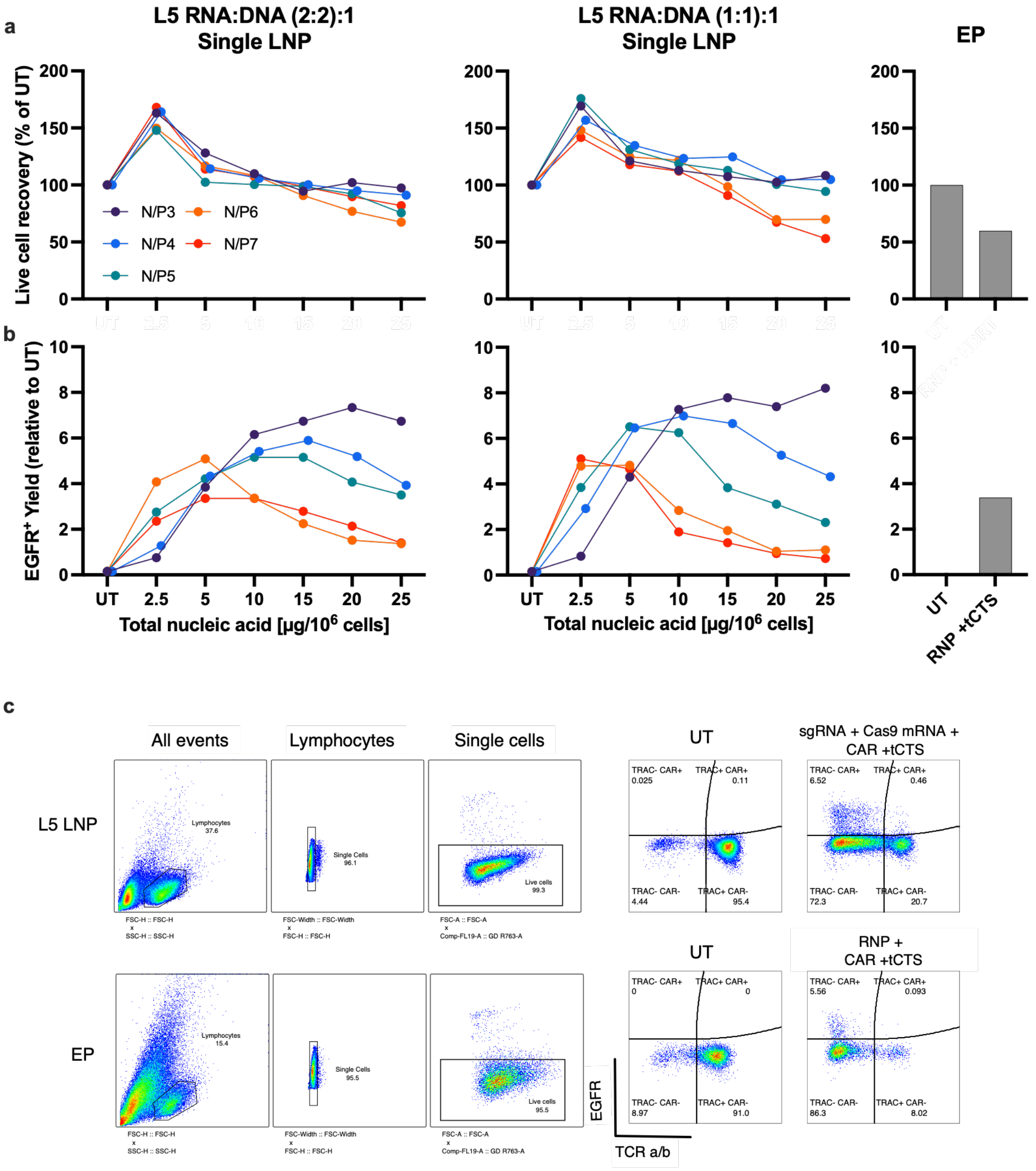


**Supplementary Figure 4. Ethanol injection formulated L5 LNP cargo and N:P ratio optimization for CAR-EGFR HDR template +tCTS delivery.** L5 LNPs co-encapsulating TRAC sgRNA, Cas9 mRNA, and CAR-EGFR HDR template +tCTS were formulated by ethanol injection and evaluated across multiple nucleic acid doses and two RNA:DNA ratios. Graphs are organized by (sgRNA: Cas9 mRNA): DNA ratio or delivery method: (2:2):1 L5, (1:1):1 L5, and EP delivery of RNP + HDR template. Day 3 flow cytometry was used to assess **(a)** live cell recovery, expressed as a percentage of UT, and **(b)** EGFR+ cell yield, calculated as the product of relative live cell recovery (normalized to UT) and $EGFR^{+}$ frequency. **(c)** Representative day 3 flow cytometry plots. Same UT wells were used for each NP curve. UT, untreated; tCTS, truncated Cas9 target sequence; *N* = 1 biological donor.

a

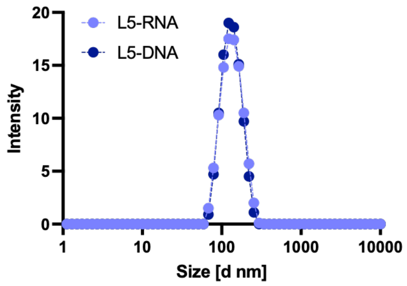


b

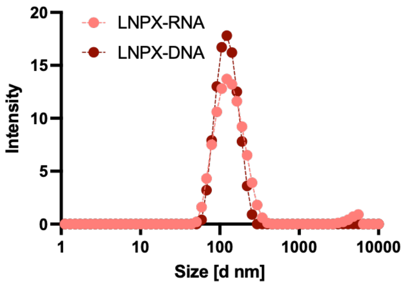


c

| Sample | Z-Ave | PDI |
|---|---|---|
| L5-RNA | 127.5 | 0.071 |
| L5-DNA | 126.4 | 0.064 |
| LNPX-RNA | 128 | 0.197 |
| LNPX-DNA | 117.4 | 0.088 |

d

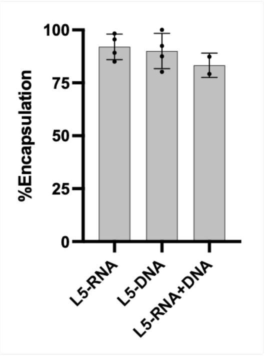


e

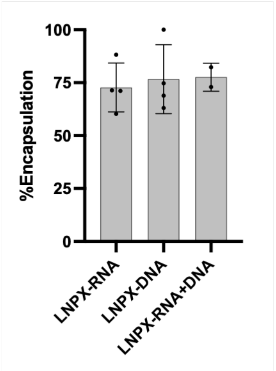


f

| Sample | %Encapsulation Efficiency | +/- SD |
|---|---|---|
| L5-RNA | 92.00 | 6.00 |
| L5-DNA | 90.04 | 8.34 |
| L5-RNA+DNA | 83.29 | 5.78 |
| LNPX-RNA | 72.73 | 11.53 |
| LNPX-DNA | 76.65 | 16.28 |
| LNPX-RNA+DNA | 77.59 | 6.60 |

**Supplementary Figure 5. Size distribution and encapsulation efficiency of NanoAssemblr™ Spark™ mixed L5 and LNPX LNP formulations. (a-c)** Representative DLS intensity distributions showing the hydrodynamic diameter of dual-encapsulated L5 and LNPX. **(d-f)** Encapsulation efficiency of single- and dual-encapsulated RNA and RNA/DNA LNP formulations, quantified by RiboGreen assay.

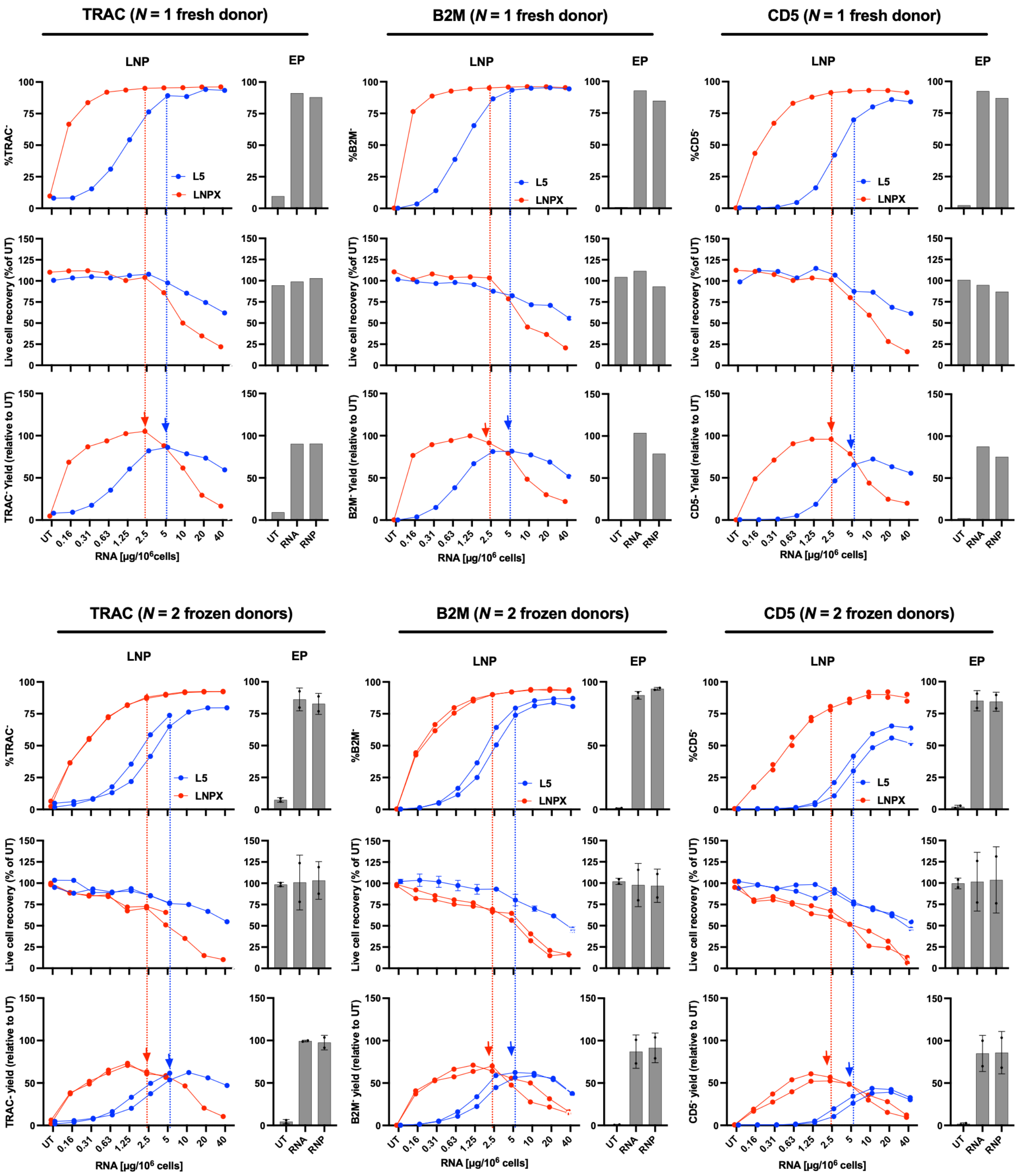
TRAC (N = 1 fresh donor)
B2M (N = 1 fresh donor)
CD5 (N = 1 fresh donor)
a
b
c
LNP
EP
%TRAC-
%B2M-
%CD5-
L5
LNPX
Live cell recovery (%of UT)
TRAC- Yield (relative to UT)
B2M- Yield (relative to UT)
CD5- Yield (relative to UT)
UT
0.16
0.31
0.63
1.25
2.5
5
10
20
40
RNA
RNP
RNA [μg/10^6 cells]
TRAC (N = 2 frozen donors)
B2M (N = 2 frozen donors)
CD5 (N = 2 frozen donors)
d
e
f
Live cell recovery (% of UT)
TRAC- yield (relative to UT)
B2M- yield (relative to UT)
CD5- yield (relative to UT)

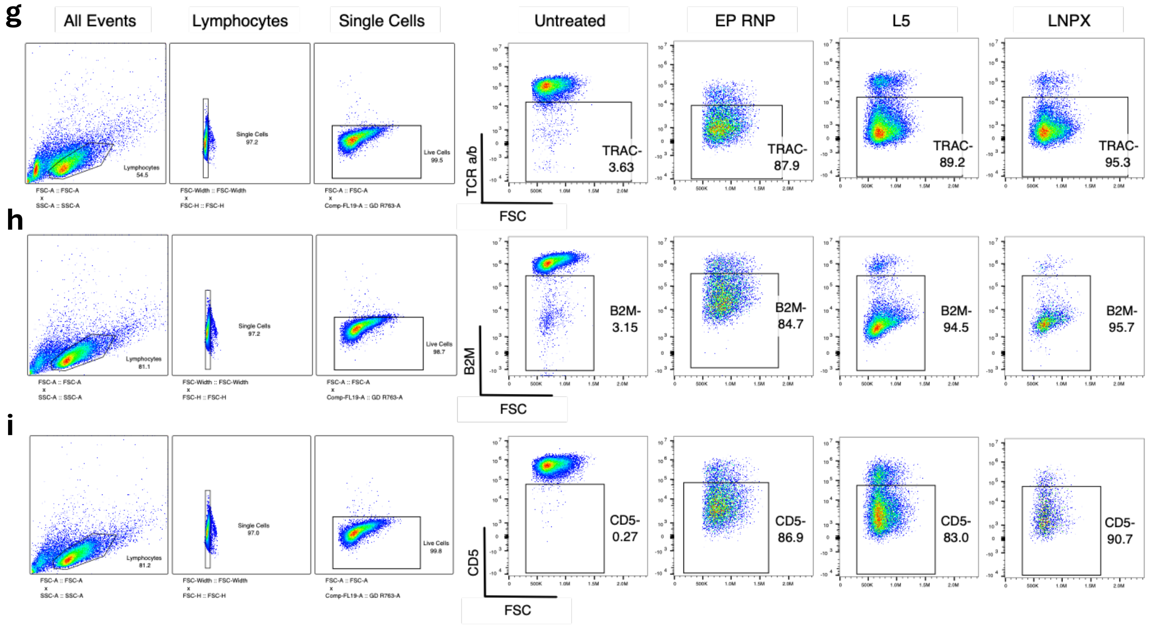


**Supplementary Figure 6. L5 and LNPX delivery of Cas9 mRNA and sgRNA targeting TRAC, B2M, or CD5 enables efficient, dose-dependent nuclear editing.** L5 and LNPX LNPs co-encapsulating Cas9 mRNA and sgRNA targeting TRAC, B2M or CD5 were formulated using the NanoAssemblr™ Spark™. Graphs are organized by target gene with TRAC, B2M and CD5 conditions shown across columns and paired LNP and EP plots shown for each target. (**a-c)** Editing outcomes in one fresh donor. **(d-f)** Editing outcomes in two frozen donors. Day 3 flow cytometry was used to assess **(a, d)** gene disruption efficiency, shown as the percentage of target negative cells, **(b, e)** live cell yield, expressed as a percentage of UT, and **(c, f)** edited cell yield, calculated as the product of relative live cell recovery (normalized to UT) and edited cell frequency. Representative day 3 flow cytometry plots for **(g)** TRAC, **(h)** B2, **(i)** CD5. UT, untreated; *N* = 1 biological donor.

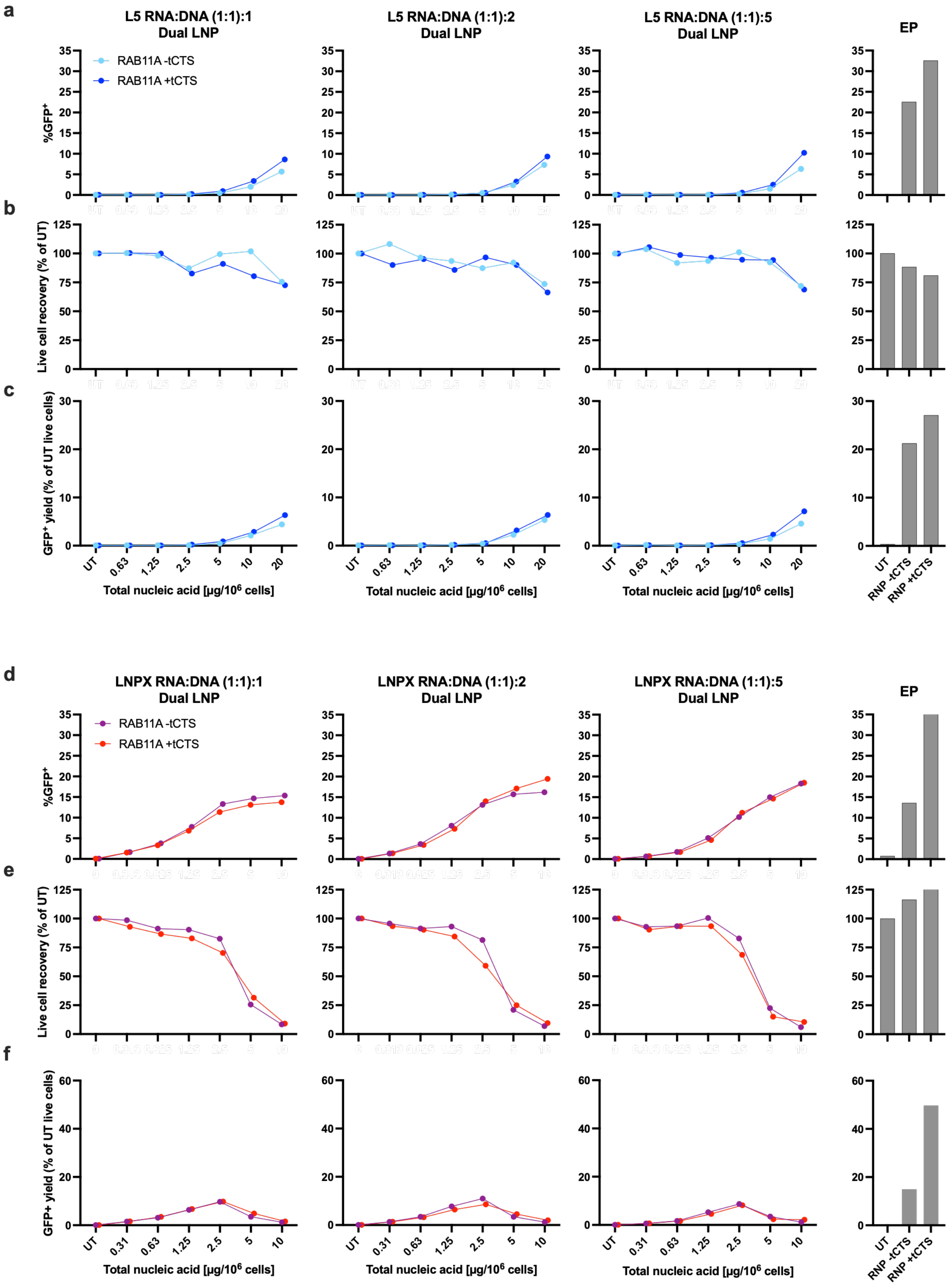
a
L5 RNA:DNA (1:1):1 Dual LNP
L5 RNA:DNA (1:1):2 Dual LNP
L5 RNA:DNA (1:1):5 Dual LNP
EP
RAB11A -tCTS
RAB11A +tCTS
%GFP+
b
Live cell recovery (% of UT)
c
GFP+ yield (% of UT live cells)
UT
0.63
1.25
2.5
5
10
20
Total nucleic acid [µg/10^6 cells]
RNP -tCTS
RNP +tCTS
d
LNPX RNA:DNA (1:1):1 Dual LNP
LNPX RNA:DNA (1:1):2 Dual LNP
LNPX RNA:DNA (1:1):5 Dual LNP
e
f
0.31

g

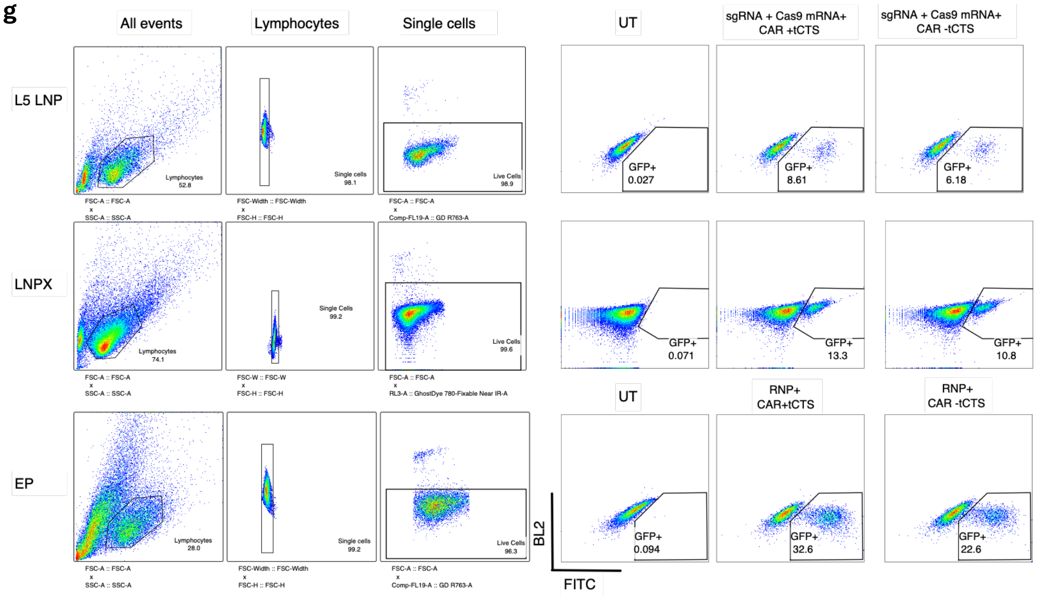


**Supplementary Figure 7. L5 and LNPX enable delivery of Cas9 mRNA, RAB11A sgRNA, and RAB11A-GFP HDR template.** L5 and LNPX LNPs were formulated using the NanoAssemblr™ Spark™ to separately encapsulate RNA (RAB11A sgRNA, Cas9 mRNA) and DNA (RAB11A-GFP HDR template +/- tCTS) cargoes. Graphs are organized by (sgRNA: Cas9mRNA) :DNA ratio or delivery method: (1:1):1, (1:1):2, (1:1):5 and EP controls delivery RNP + HDR template +/- tCTS. (**a-c)** L5 LNP-mediated delivery. **(d-f)** LNPX LNP-mediated delivery. Day 3 flow cytometry was used to assess **(a, d)** editing efficiency, shown as the percentage of GFP$^{+}$ cells, **(b, e)** live cell yield, expressed as a percentage of UT, and **(c, f)** edited cell yield, calculated as the product of relative live cell recovery (normalized to UT) and GFP$^{+}$ frequency. Representative day 3 flow cytometry plots for **(g)** L5 and **(h)** LNPX. UT, untreated; tCTS, truncated Cas9 target sequences; *N* = 1 biological donor.

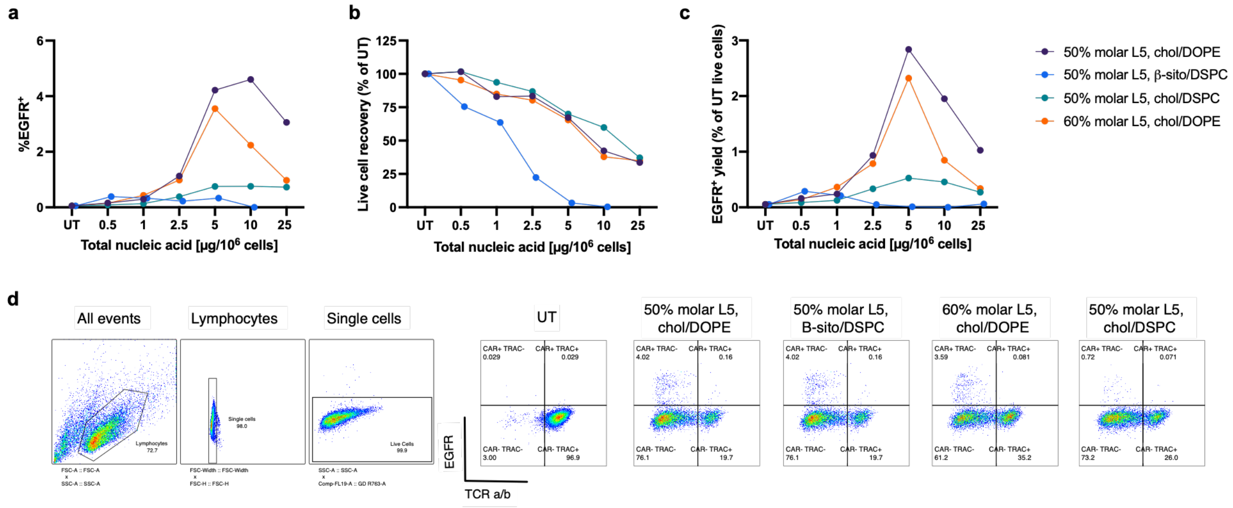


**Supplementary Figure 8. Varying L5 formulation parameters to evaluate knock-in efficacy for CAR-EGFR HDR template +tCTS.** L5 LNPs co-encapsulating TRAC sgRNA, Cas9 mRNA, and CAR-EGFR HDR template were formulated by ethanol injection with varying molar percentages of L5 ionizable lipid, 50% or 60%, and different helper lipid and sterol compositions, including DOPE or DSPC and cholesterol or β-sitosterol. Day 3 flow cytometry was used to assess **(a)** editing efficiency, shown as percentage of $EGFR^+$ cells, **(b)** live cell yield, expressed as a percentage of UT, and (**c**) edited cell yield, calculated as the product of relative live cell recovery (normalized to UT) and $EGFR^+$ frequency. **(d)** Representative flow cytometry plots at day 3 for each LNP and EP condition. UT, untreated; tCTS, truncated Cas9 target sequences; *N* = 1 biological donor.

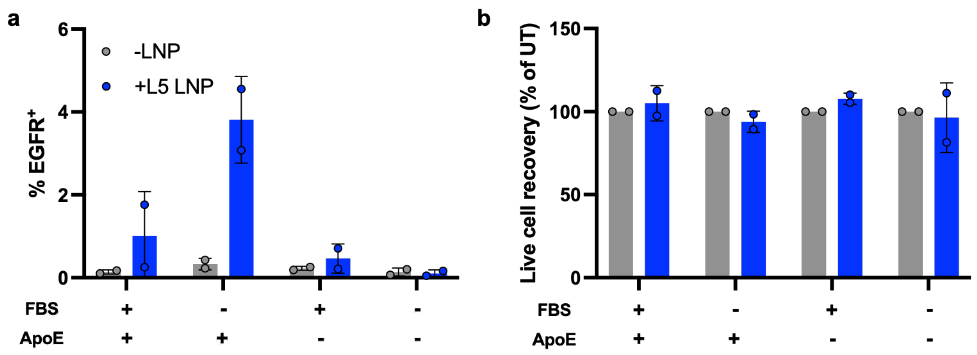


**Supplementary Figure 9. ApoE supplementation is required for efficient L5 LNP-mediated knock-in.** L5 LNPs co-encapsulating TRAC sgRNA, Cas9 mRNA, and CAR-EGFR HDR template +tCTS were formulated by ethanol injection 1:1:1 w/w/w ratio. T cells were treated with dose of 15 µg nucleic acid/$10^6$ cells and subsequently cultured in media with or without 5% FBS and with or without 1 ug/mL ApoE. Day 3 flow cytometry was used to assess **(a)** editing efficiency, shown as percentage of EGFR$^+$ cells, **(b)** live cell yield, expressed as a percentage of UT. UT, untreated; ApoE = apolipoprotein E; tCTS, truncated Cas9 target sequences; *N* = 1 biological donor.

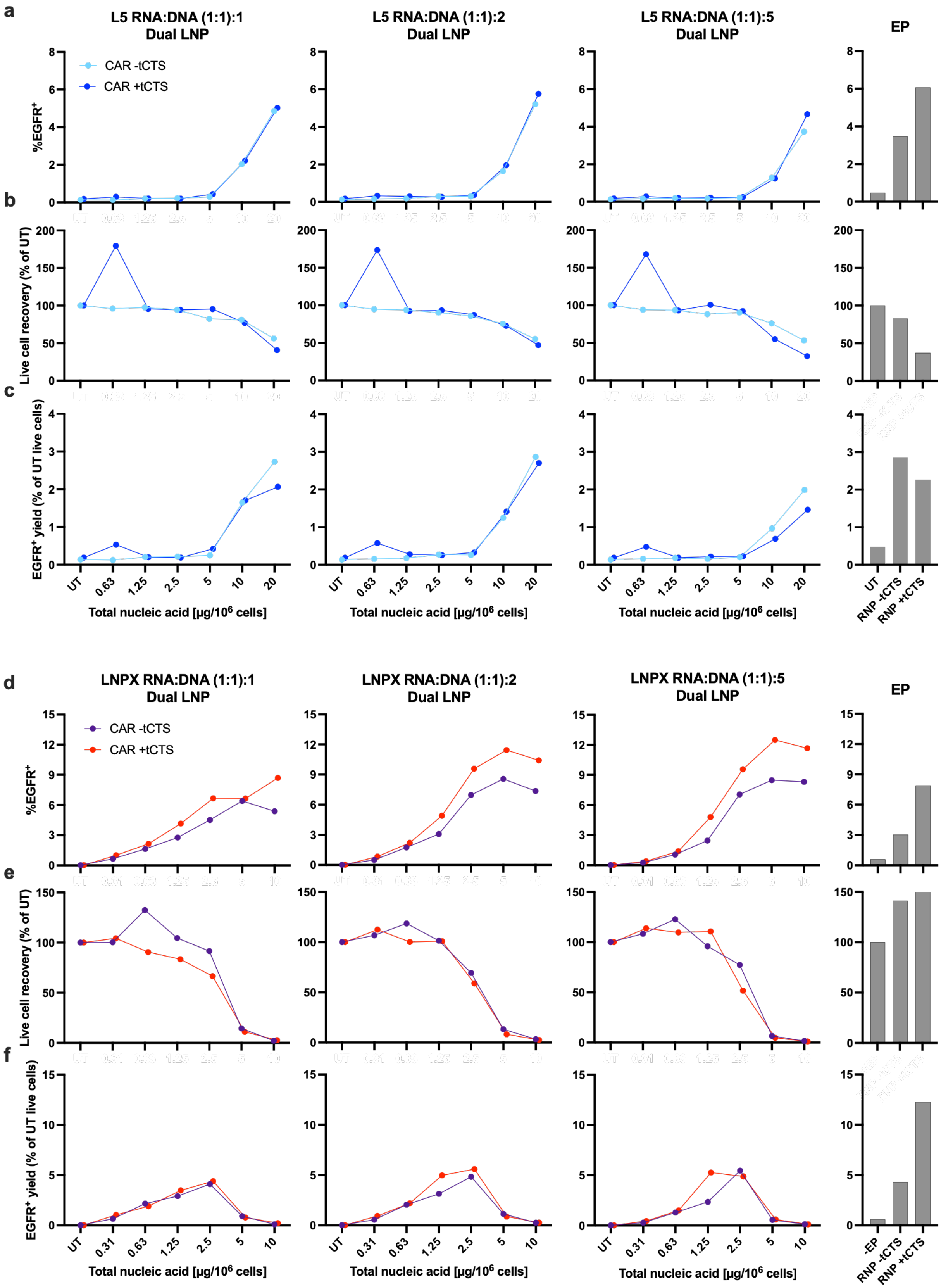
a
L5 RNA:DNA (1:1):1 Dual LNP
L5 RNA:DNA (1:1):2 Dual LNP
L5 RNA:DNA (1:1):5 Dual LNP
EP
CAR -tCTS
CAR +tCTS
%EGFR+
b
Live cell recovery (% of UT)
c
EGFR+ yield (% of UT live cells)
UT
0.63
1.25
2.5
5
10
20
Total nucleic acid [µg/10^6 cells]
RNP -tCTS
RNP +tCTS
d
LNPX RNA:DNA (1:1):1 Dual LNP
LNPX RNA:DNA (1:1):2 Dual LNP
LNPX RNA:DNA (1:1):5 Dual LNP
EP
CAR -tCTS
CAR +tCTS
%EGFR+
e
Live cell recovery (% of UT)
f
EGFR+ yield (% of UT live cells)
UT
0.31
0.63
1.25
2.5
5
10
Total nucleic acid [µg/10^6 cells]
-EP
RNP -tCTS
RNP +tCTS

**g**

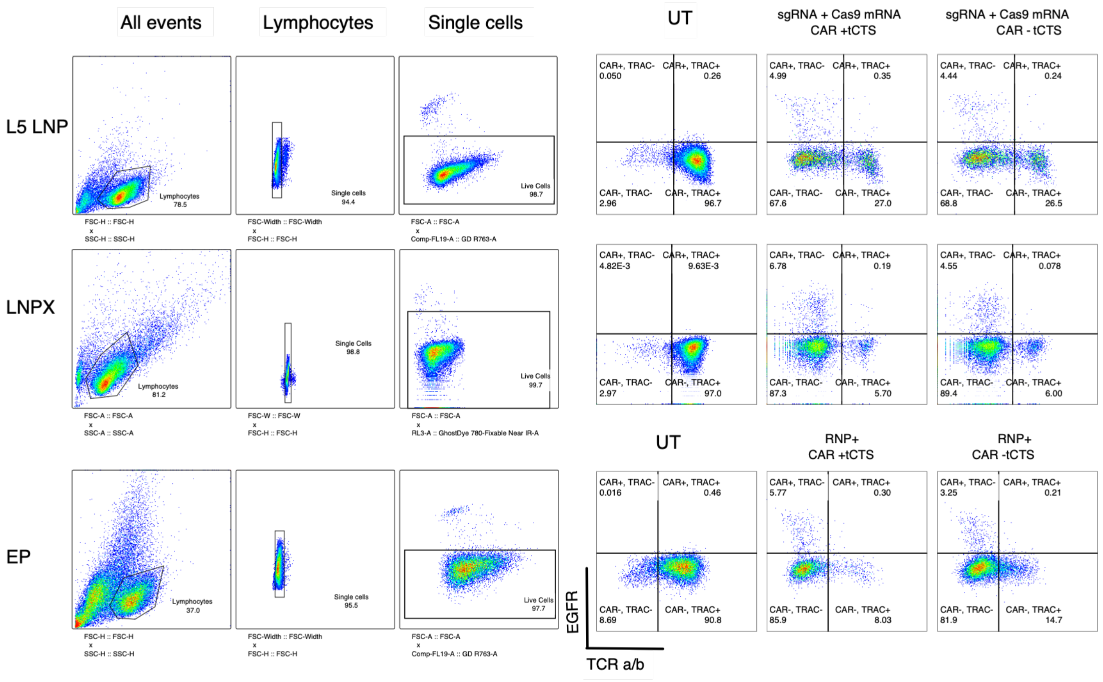


**Supplementary Figure 10. L5 and LNPX enables delivery of Cas9 mRNA, TRAC sgRNA, and CAR-EGFR HDR template.** L5 and LNPX LNPs separately encapsulating RNA (TRAC sgRNA, Cas9 mRNA) and DNA (CAR-EGFR HDR template +/- tCTS) cargoes were formulated using the NanoAssemblr™ Spark™. **(a-c)** Graphs are organized by RNA:DNA ratio or delivery method: (1:1):1, (1:1):2, (1:1):5 and electroporation controls delivery RNP + HDR template +/- tCTS. (**a-c)** L5 LNP-mediated delivery. **(d-f)** LNPX LNP-mediated delivery. Day 3 flow cytometry was used to assess **(a, d)** editing efficiency, shown as the percentage of EGFR$^+$ cells, **(b, e)** live cell yield, expressed as a percentage of UT, and **(c, f)** edited cell yield, calculated as the product of relative live cell recovery (normalized to UT) and EGFR$^+$ frequency. **(g)** Representative day 3 flow cytometry plots for L5, LNPX, and EP samples. UT, untreated; tCTS, truncated Cas9 target sequences; *N* = 1 biological donor.

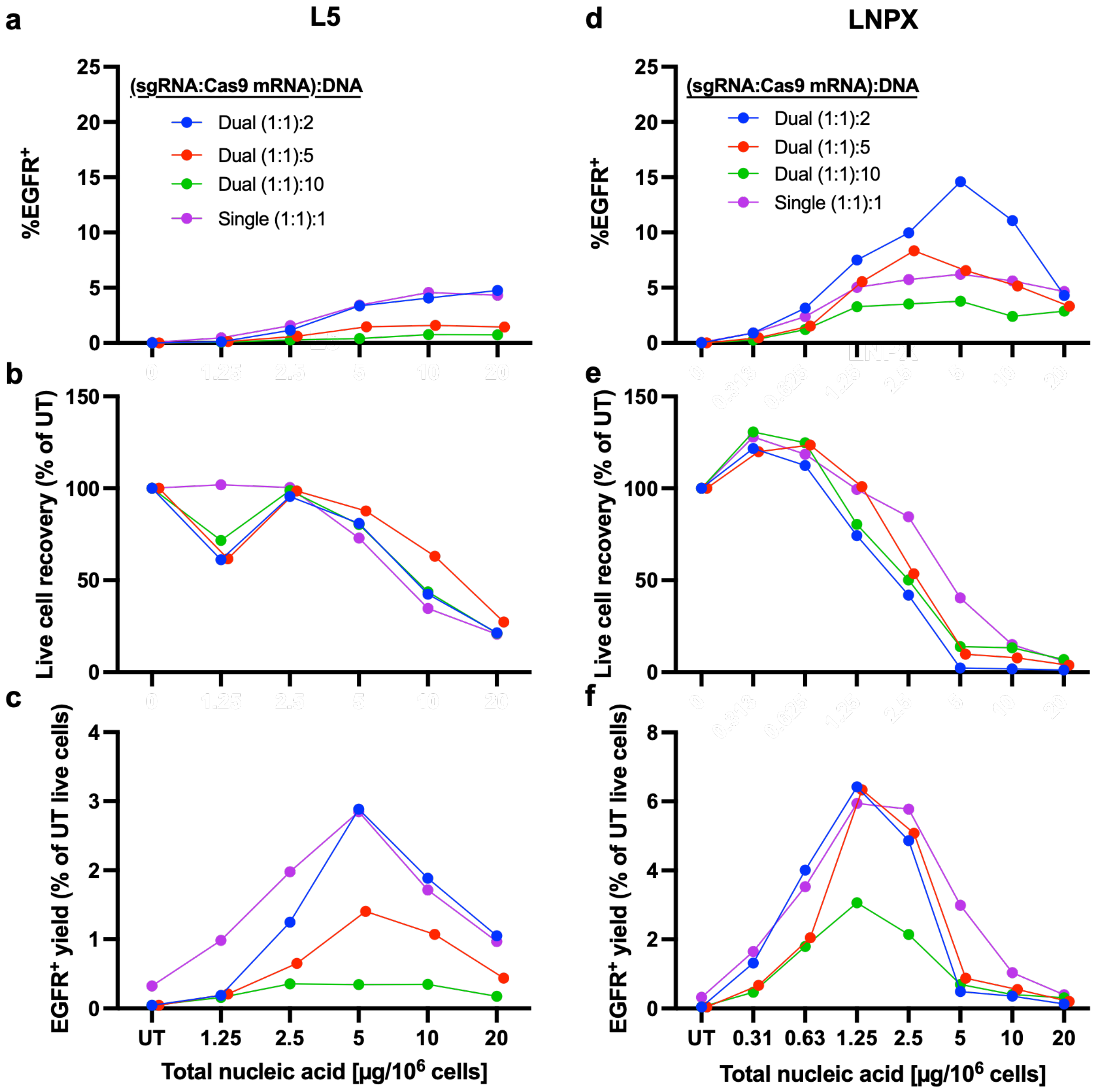


**Supplementary Figure 11: Dual- and single- encapsulated LNPs achieve similar CAR-EGFR knock-in yields.** L5 and LNPX LNPs were formulated either by co-encapsulating sgRNA, Cas9 mRNA, and CAR-EGFR HDR template +tCTS or by separately encapsulating the RNA and the DNA components using the NanoAssemblr™ Spark™. Dual encapsulated LNPs had (sgRNA:cas9 mRNA):DNA w/w ratio of (1:1):2, (1:1):5, (1:1):10. Single encapsulated LNPs are formulated at a (sgRNA:cas9 mRNA):DNA w/w ratio of (1:1):1. Graphs are organized by LNP formulation with L5 shown on the left and LNPX shown on the right. Day 3 flow cytometry was used to assess **(a, d)** editing efficiency, shown as the percentage of EGFR$^{+}$ cells, **(b, e)** live cell yield, expressed as a percentage of UT, and **(c, f)** edited cell yield, calculated as the product of relative live cell recovery (normalized to UT) and EGFR$^{+}$ frequency. UT, untreated; tCTS, truncated Cas9 target sequences; *N* = 1 biological donor.

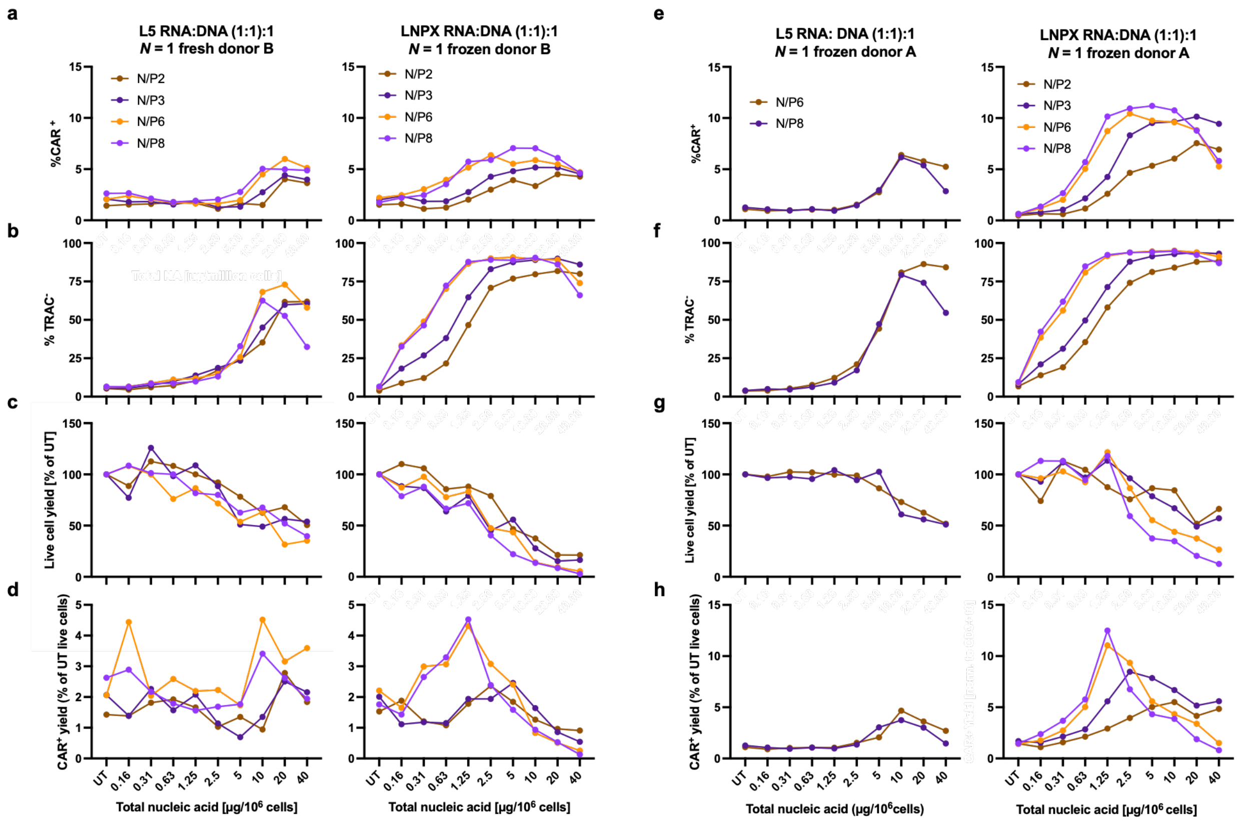


**Supplementary Figure 12. N/P ratio optimization on LNP-mediated HDR editing for CAR-EGFR HDR template knock-in.** L5 and LNPX LNPs encapsulating TRAC sgRNA, Cas9 mRNA, and a CAR-EGFR HDR template +tCTS at a 1:1:1 weight ratio were formulated using the NanoAssemblr™ Spark™ and evaluated across three independent donors, including one donor shown in Figure 2c-d. Graphs are organized by donor with **(a-d)** one fresh donor and **(e-g)** one frozen donor shown showing LNP conditions and corresponding EP controls delivery RNA or RNP + HDR template + tCTS. Day 3 flow cytometry was used to assess **(a, e)** editing efficiency, shown as the percentage of $EGFR^{+}$ cells, **(b, f)** TRAC disruption, shown as a percentage of $TRAC^{-}$ cells, **(c,g)** live cell yield, expressed as a percentage of UT, and **(d, h)** edited cell yield, calculated as the product of relative live cell recovery (normalized to UT) and $CAR^{+}$ frequency. UT, untreated; tCTS, truncated Cas9 target sequences; *N* = 2 biological donors.

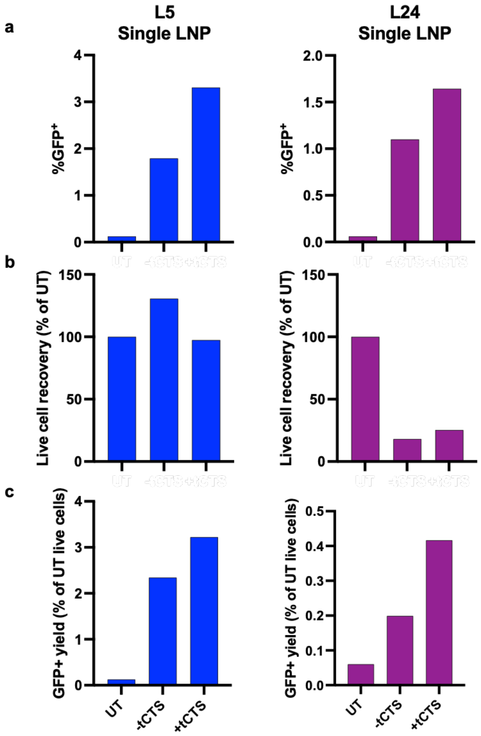


**Supplementary Figure 13. Addition of tCTS enhances RAB11A-GFP knock-in in L24 and L5 LNP systems.** L5 and L24 LNPs co-encapsulating RAB11A sgRNA, Cas9 mRNA, and a RAB11A-GFP HDR template +/- tCTS were formulated by ethanol injection. Day 3 flow cytometry was used to assess **(a)** editing efficiency, shown as percentage of $GFP^+$ cells, **(b)** live cell yield, expressed as a percentage of UT, and **(c)** edited cell yield, calculated as the product of relative live cell recovery (normalized to UT) and $GFP^+$ frequency. UT, untreated; tCTS, truncated Cas9 target sequences; *N* = 1 biological donor.

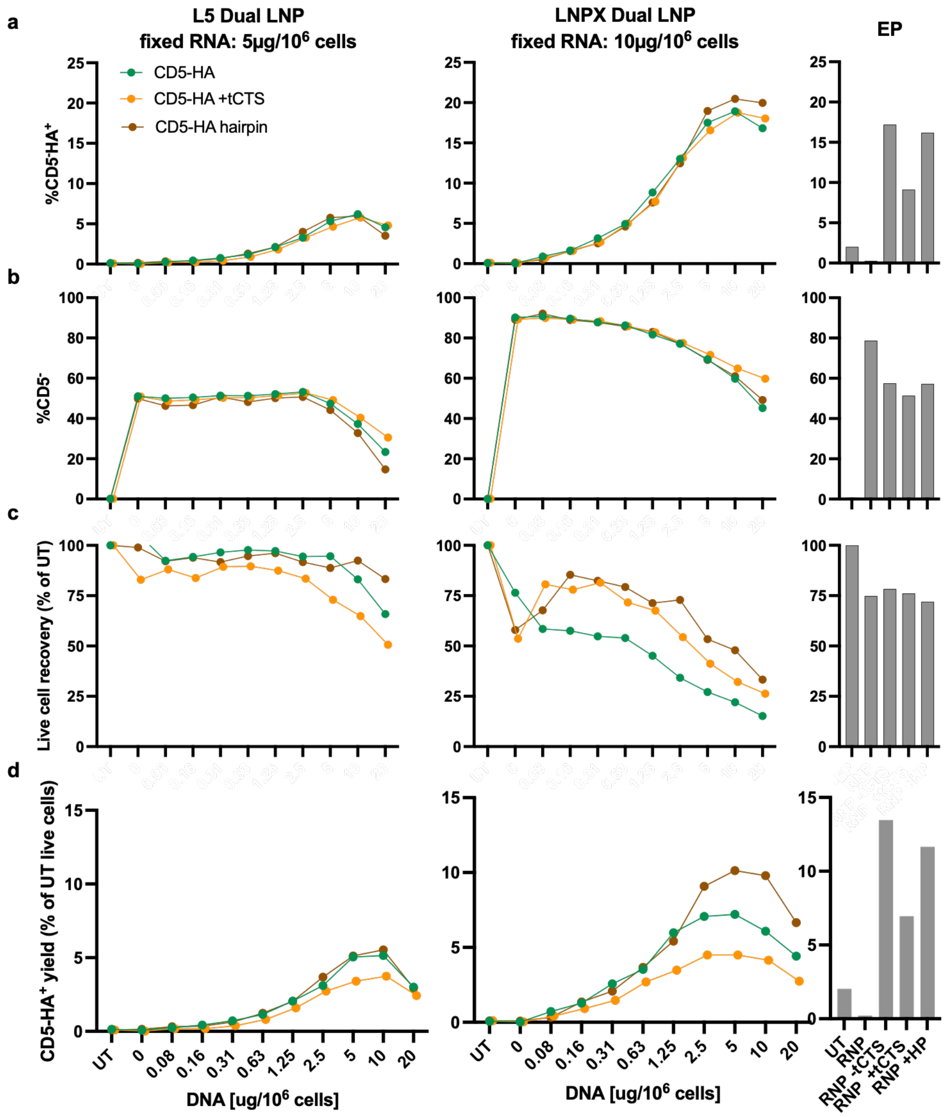


**Supplementary Figure 14. L5 and LNPX delivery of CD5-HA ssODN.** L5 and LNPX LNPs separately encapsulating RNA (TRAC sgRNA, Cas9 mRNA) and DNA (CAR-EGFR HDR template +/- tCTS) cargoes were formulated using the NanoAssemblr™ Spark™. L5 conditions were treated at a fixed RNA dose of 5µg/10$^6$ cells and LNPX conditions were treated at a fixed RNA dose of 2.5µg/10$^6$ cells, both with a dose range for the DNA LNP. Day 3 flow cytometry was used to assess **(a)** editing efficiency, shown as the percentage of CD5-HA$^+$ cells, **(b)** CD5 disruption, shown as a percentage of CD5$^-$ cells, **(c)** live cell yield, expressed as a percentage of UT, and **(d)** edited cell yield, calculated as the product of relative live cell recovery (normalized to UT) and CD5-HA$^+$ frequency. UT, untreated; tCTS, truncated Cas9 target sequences; *N* = 1 biological donor.

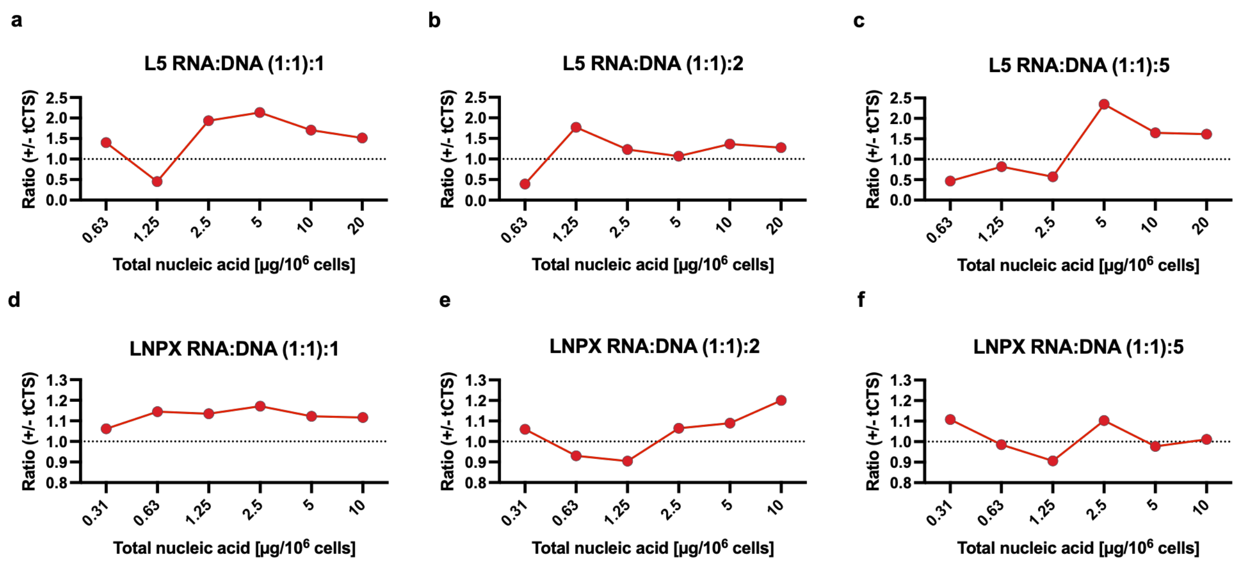


**Supplementary Figure 15. Comparison of tCTS versus no tCTS addition to RAB11A-GFP HDR template in L5 and LNPX LNPs.** L5 and LNPX LNPs separately encapsulating RNA (RAB11A sgRNA andCas9 mRNA) and DNA (RAB11A-GFP HDR template +/- tCTS) cargoes were formulated using the NanoAssemblr™ Spark™. LNPs were treated at 3 different (sgRNA:Cas9 mRNA):DNA ratios. Day 3 flow cytometry was used to assess editing efficiency and plots display the +tCTS/-tCTS ratio of HDR efficiency for **(a-c)** L5 and **(d-f)** LNPX. Ratios >1 indicate improved HDR with the addition of tCTS. tCTS, truncated Cas9 target sequences; *N* = 1 biological donor.

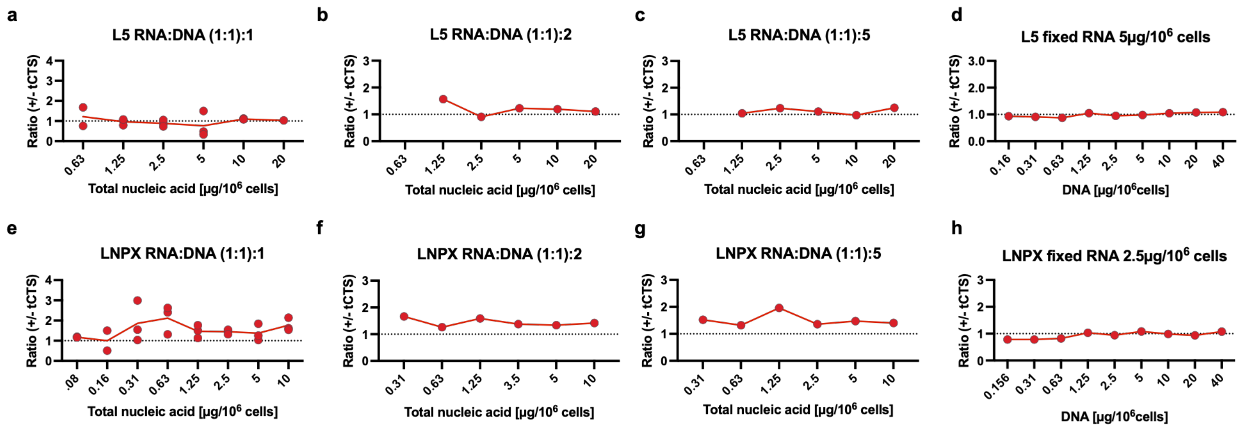


**Supplementary Figure 16. Comparison of tCTS versus no tCTS addition to CAR-EGFR HDR template in L5 and LNPX LNPs.** L5 and LNPX LNPs separately encapsulating RNA (TRAC sgRNA andCas9 mRNA) and DNA (TRAC-EGFR HDR template +/- tCTS) cargoes were formulated using the NanoAssemblr™ Spark™. LNPs were treated at either fixed dose **(a-c, e-g)** or fixed RNA dose **(d, h)**. Day 3 flow cytometry was used to assess editing and plots display the +tCTS/-tCTS ratio of HDR efficiency for **(a-d)** L5 and **(e-h)** LNPX. Ratios >1 indicate improved HDR with the addition of tCTS. tCTS, truncated Cas9 target sequences; *N* = 1 biological donor.

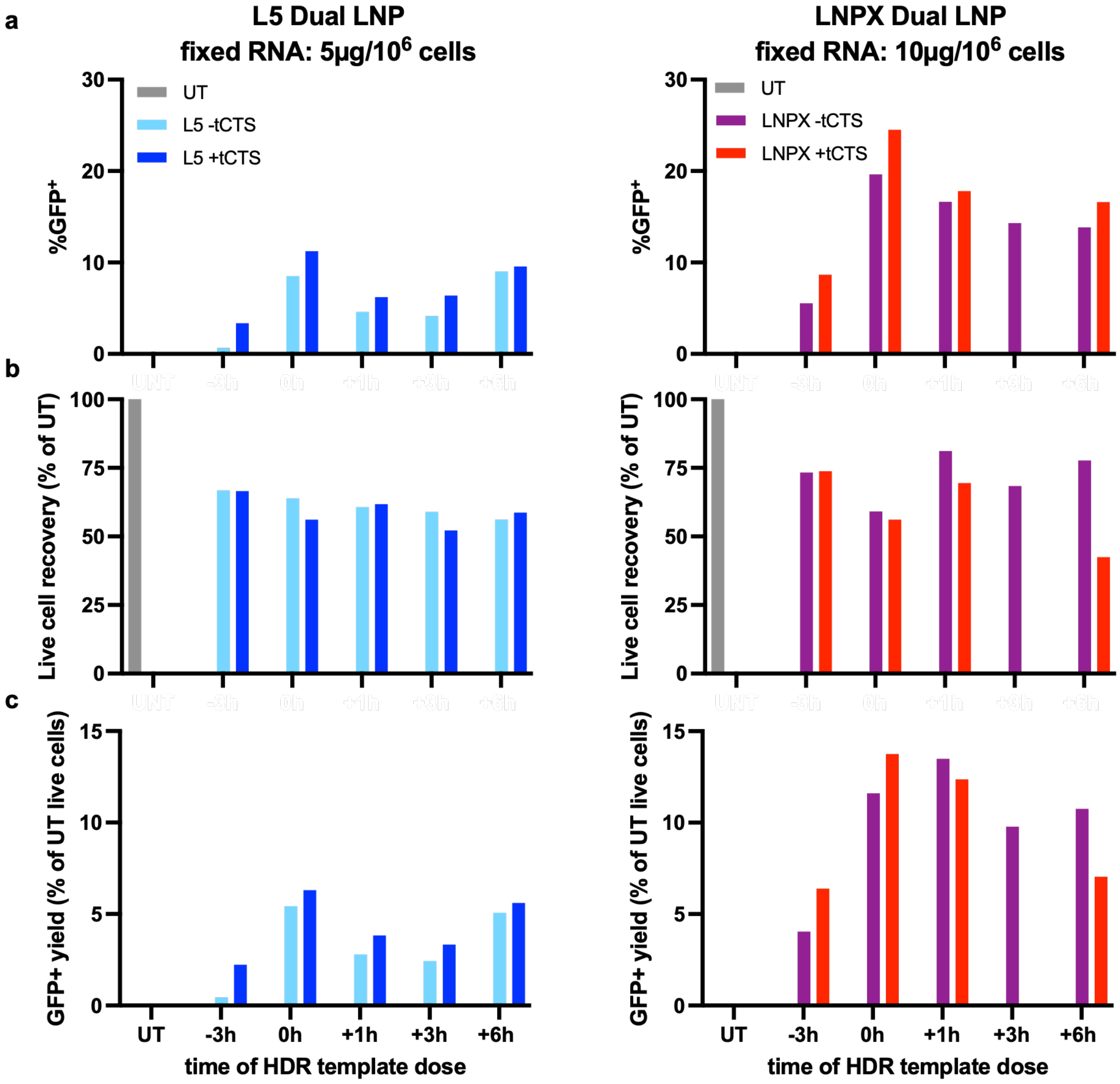


**Supplementary Figure 17. Highest RAB11A-GFP transfection rates occur with simultaneous dosing of separately encapsulated RNA and HDR template LNPs.** L5 and LNPX LNPs separately encapsulating RNA (RAB11A sgRNA andCas9 mRNA) and DNA (RAB11A-GFP HDR template +/- tCTS) cargoes were formulated using the NanoAssemblr™ Spark™. L5 RNA LNPs were treated at 10 µg nucleic acid/$10^6$ cells with (1:1):2 (sgRNA:cas9 mRNA):HDRT ratios and LNPX RNA LNPs treated at 2.5 µg nucleic acid/$10^6$ cells, (1:1):2 (sgRNA:cas9 mRNA):HDRT ratios at t=0. HDRT was added at various time points. Day 3 flow cytometry was used to assess **(a)** editing efficiency, shown as the percentage of $GFP^+$ cells, **(b)** live cell yield, expressed as a percentage of UT, and **(c)** edited cell yield, calculated as the product of relative live cell recovery (normalized to UT) and $GFP^+$ frequency. UT, untreated; tCTS, truncated Cas9 target sequences; *N* = 1 biological donor.

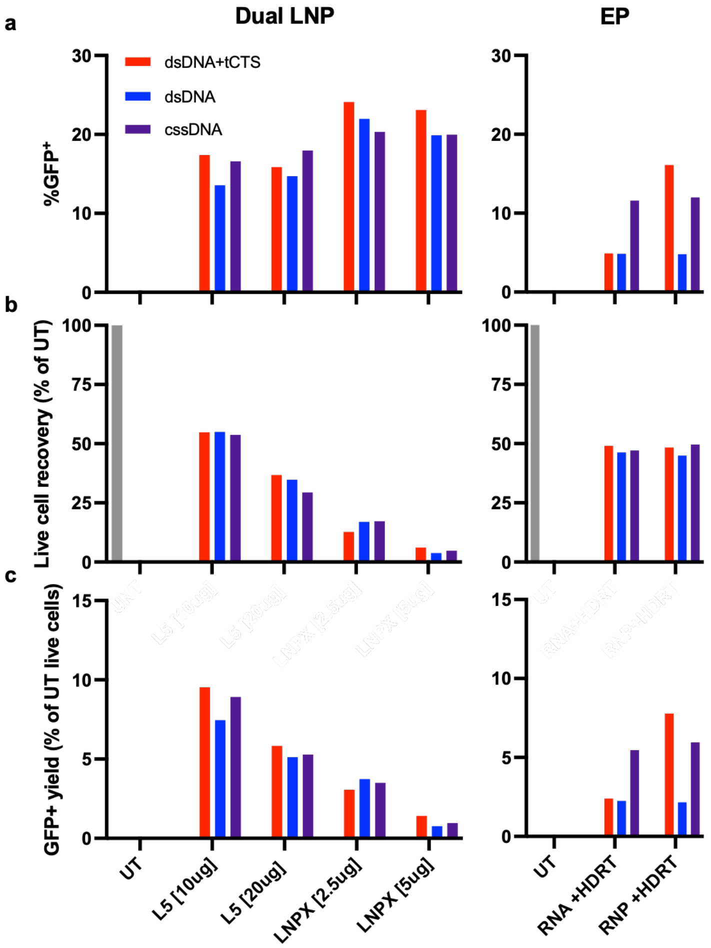


**Supplementary Figure 18: L5 and LNPX delivery of RAB11A-GFP cssDNA shows comparable KI to dsDNA delivery.** LNPX LNPs encapsulating TRAC sgRNA, Cas9 mRNA, and a CAR-EGFR HDR template as dsDNA (+/-tCTS) or cssDNA were formulated using the NanoAssemblr™ Spark™. L5 RNA LNPs were treated at 10 or 20 µg nucleic acid/$10^6$ cells with (1:1):2 (sgRNA:cas9 mRNA):HDRT ratios and LNPX RNA LNPs treated at 2.5 or 5 µg nucleic acid/$10^6$ cells, (1:1):2 (sgRNA:cas9 mRNA):HDRT ratios. HDRT was added at various time points. were treated at a fixed RNA dose of 2.5µg/$10^6$ cells, both with DNA doses listed. Graphs are organized by LNP and EP. Day 3 flow cytometry was used to assess **(a)** editing efficiency, shown as the percentage of $GFP^+$ cells, **(b)** live cell yield, expressed as a percentage of UT, and **(c)** edited cell yield, calculated as the product of relative live cell recovery (normalized to UT) and $GFP^+$ frequency. UT, untreated; tCTS, truncated Cas9 target sequences; *N* = 1 biological donor.

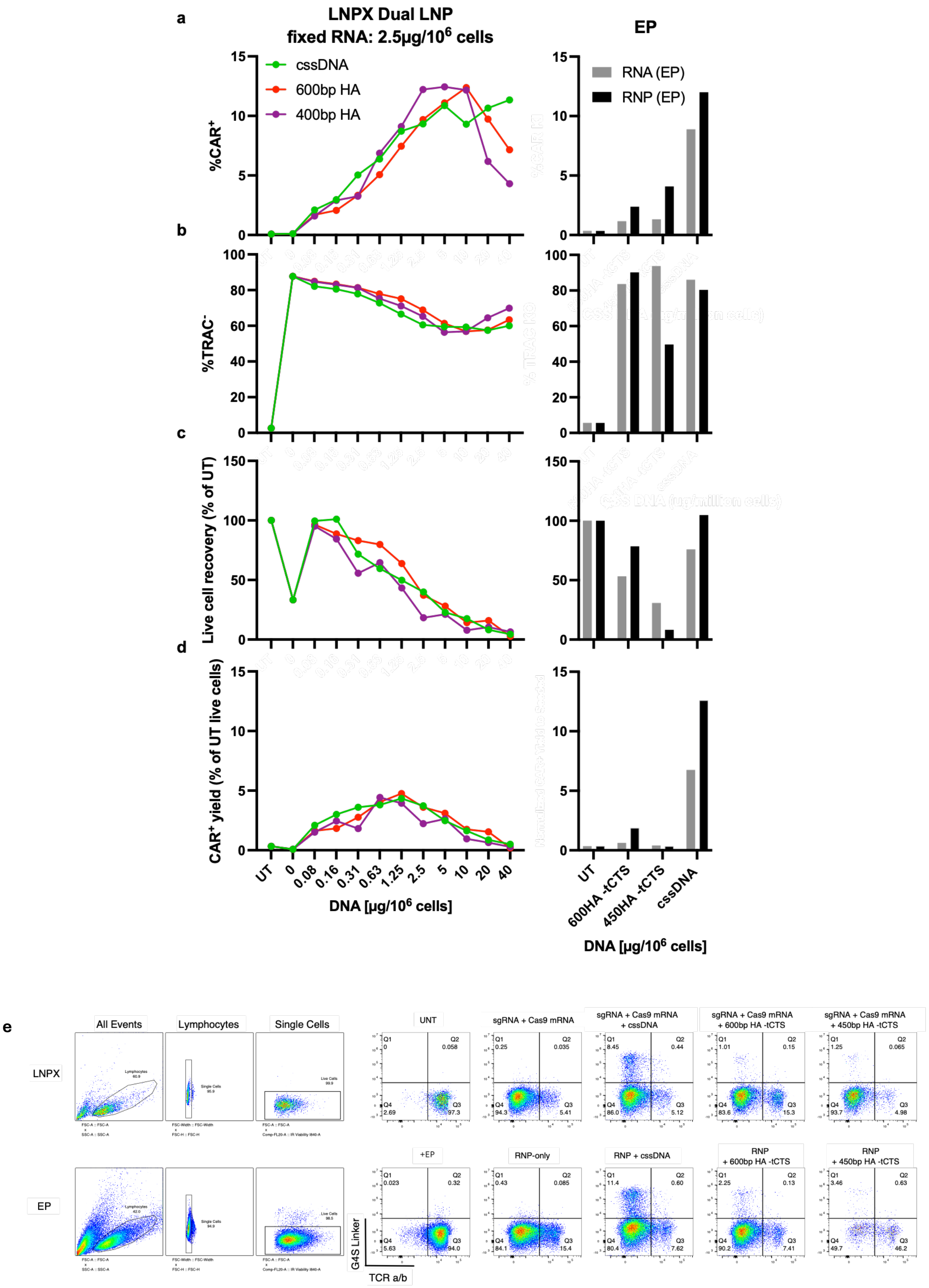


**Supplementary Figure 19: Editing efficiency of CAR cssDNA and CAR-EGFR HDR template 600bp versus 450bp homology arms on each side.** LNPX LNPs encapsulating TRAC sgRNA, Cas9 mRNA, or a CAR template (cssDNA 300bp HA with no EGFR, 400bp HA or 600bp HA CAR-EGFR HDR template -tCTS) were formulated using the NanoAssemblr™ Spark™. L5 conditions were treated at a fixed RNA dose of 5μg/$10^6$ cells and LNPX conditions were treated at a fixed RNA dose of 2.5μg/$10^6$ cells, both with a dose range for the DNA LNP. Graphs are organized by LNP and EP. Day 3 flow cytometry was used to assess **(a)** editing efficiency, shown as the percentage of $CAR^+$ cells, **(b)** TRAC disruption, shown as a percentage of $TRAC^-$ cells, **(c)** live cell yield, expressed as a percentage of UT, and **(d)** edited cell yield, calculated as the product of relative live cell recovery (normalized to UT) and $CAR^+$ frequency. **(e)** Representative day 3 flow cytometry plots for LNPX, and EP samples. UT, untreated; tCTS, truncated Cas9 target sequences; bp, base pair; HA, homology arm; *N* = 1 biological donor.

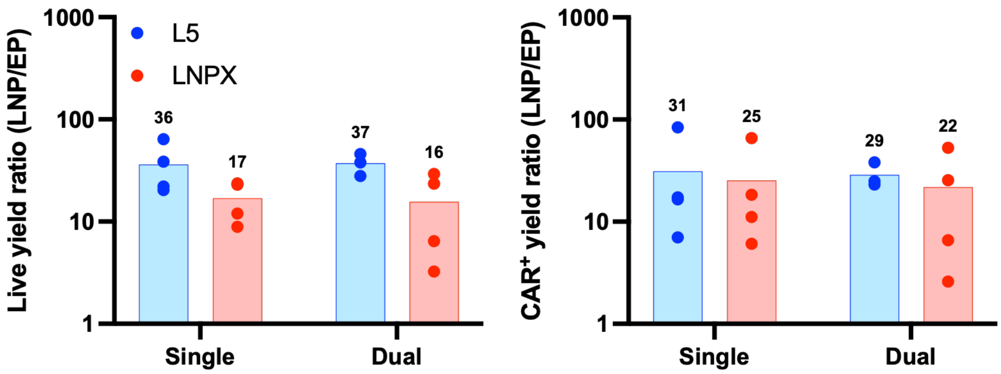


**Supplementary Figure 20. Comparison of LNP/EP ratio of live cell yields and CAR$^+$ editing yields assessed by Day 3 flow cytometry.** Ratios were calculated as the yield per input cell following LNP delivery divided by that following EP. Data are from *N* = 4 biological donors for single encapsulation and *N* = 3-4 biological donors for dual encapsulation, cultured at optimal conditions for each editing method.

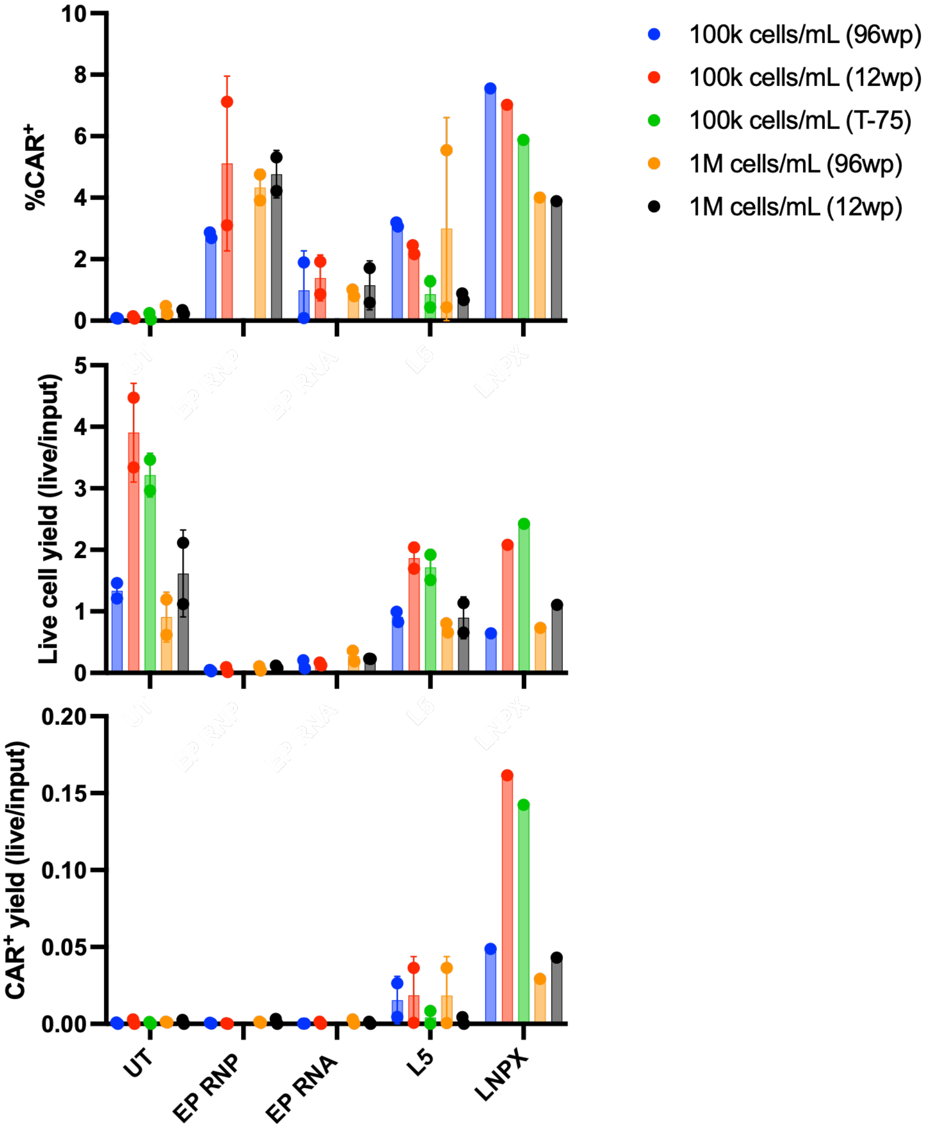


**Supplementary Figure 21: Comparison of EP and LNP editing across matched culture conditions highlights higher LNP yield compared to EP.** L5 and LNPX LNPs encapsulating TRAC sgRNA, Cas9 mRNA, and CAR-EGFR HDRT +tCTS were formulated using the NanoAssemblr™ Ignite™. L5 conditions were treated at a total dose of 10µg/$10^6$ cells and LNPX conditions were treated at a total dose of 2.5µg/$10^6$ cells. Day 3 flow cytometry was used to assess **(a)** editing efficiency, shown as the percentage of $CAR^+$ cells, **(b)** live cell yield, calculated as total live cell count divided by input cell count, and **(c)** edited $CAR^+$ cell yield, calculated as total $CAR^+$ cell count input cell count. UT, untreated; tCTS, truncated Cas9 target sequences; *N* = 1 (LNPX) and *N* = 2 (L5) biological donors across two independent experiments.

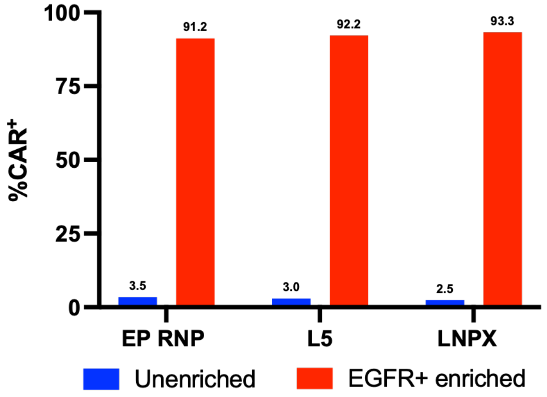


**Supplementary Figure 22. EGFR bead enrichment allows for selection of edited EGFR+ CAR T cells by EP and LNP delivery**. **(a)** Population of EGFR+ CAR T cells increased from ~3% to >90% post magnetic enrichment.

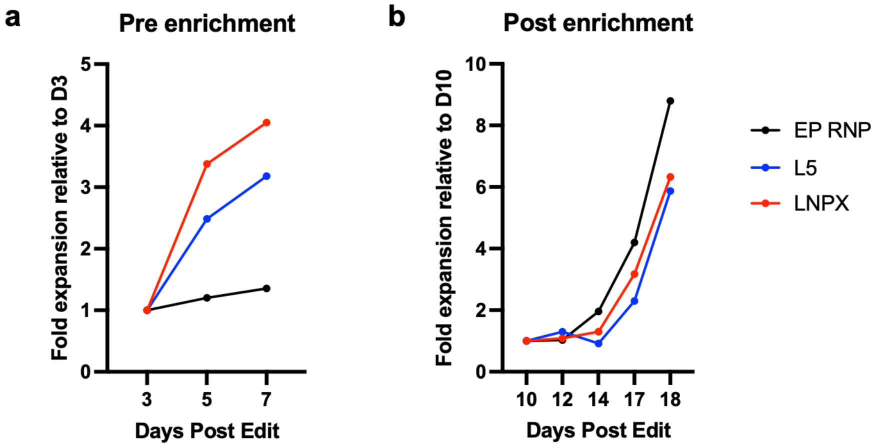


**Supplementary Figure 23. Expansion kinetics of CAR-T cells generated via LNP vs EP generated CAR-T.** Flow cytometry was used to quantify viable cell numbers over 18 days post edit. Fold expansion is shown for **(a)** the pre-enrichment phase, normalized to Day 3 cell count, and **(b)** the post-enrichment phase following EGFR magnetic enrichment, normalized to the Day 10 cell count. *N* = 1 biological donor.

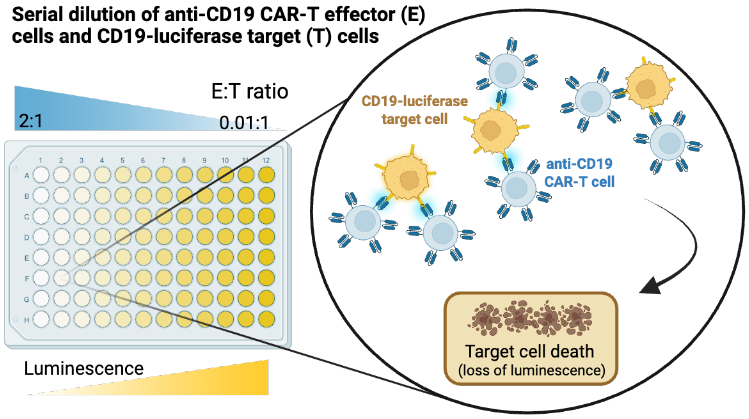


**Supplementary Figure 24. Schematic of cell killing assay**. CD19 targeted CAR T cells (E) were plated at a serial dilution from a 2:1 to 0.01:1 ratio with either CD19 NALM-6 cells or CD19 Raji cells (T) and incubated overnight to allow for targeted killing.

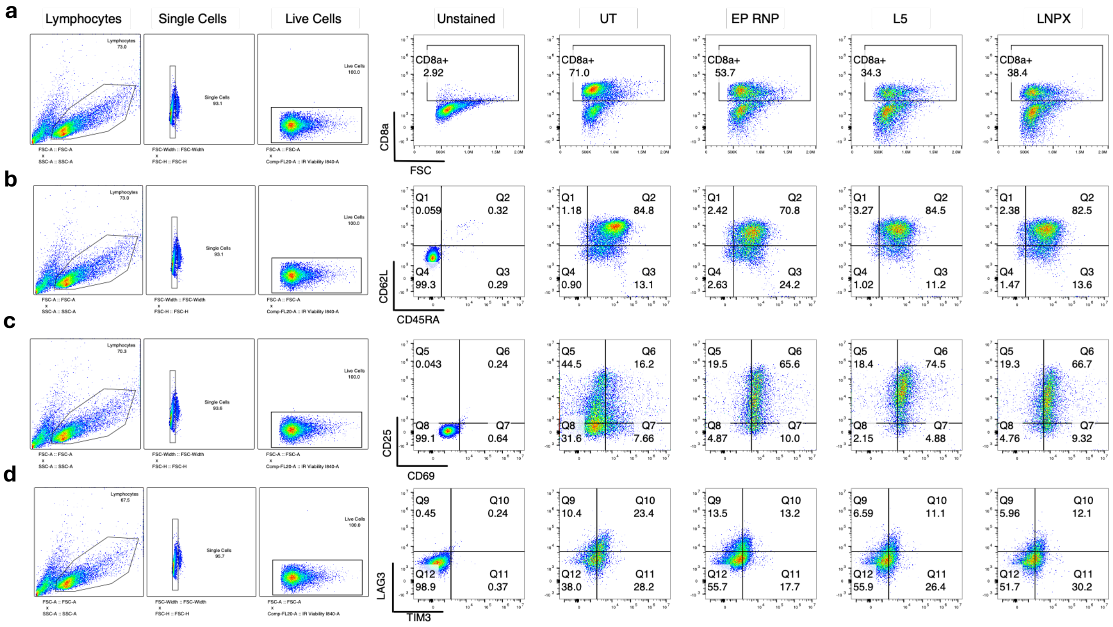


**Supplementary Figure 25. Representative flow cytometry plots for post-enrichment EGFR+ CAR T cell phenotyping displays similar phenotypes across LNP and EP generated CAR T cells.** Gating shown for **(a)** CD8a **(b)** CD62L vs CD45RA **(c)** CD25 vs CD69 and **(d)** LAG3 vs TIM3. UT= untreated; *N* = 1 biological donor.

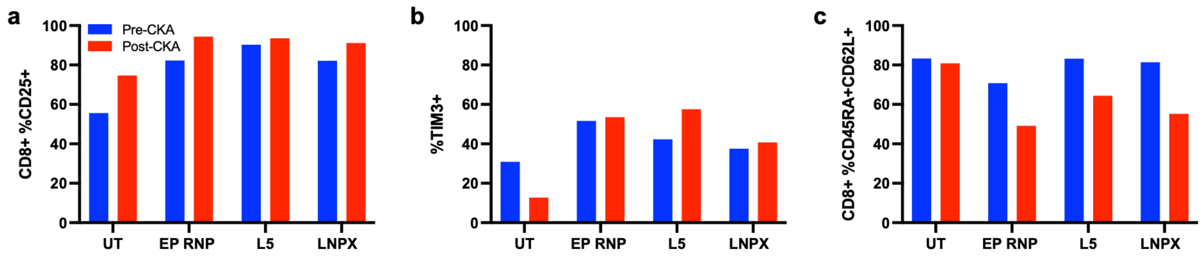


**Supplementary Figure 26. EGFR+ CAR T phenotyping before and after cell killing assay remains similar between LNP and EP generated CAR T.** Flow cytometry after 24 hours post CKA display **(a)** CD8+ CD25+ **(b)** TIM3+ and **(c)** CD8+ CD45RA+ CD62L+ levels. UT= untreated; *N* = 1 biological donor.

| Target | Fluorophore | Dilution | Manufacturer | Clone | Catalog# |
|---|---|---|---|---|---|
| B2M | FITC | 1:100 | Biolegend | 2M2 | 316304 |
| TCRa/b | PE | 1:100 | Biolegend | IP26 | 306708 |
| TCRa/b | APC | 1:100 | Biolegend | IP26 | 306718 |
| CD5 | PE | 1:100 | Biolegend | UCHT2 | 300608 |
| EGFR | AF488 | 1:100 | Biolegend | AY13 | 352908 |
| CAR GS-Linker | PE | 1:100 | Cell Signaling Tech | E702V | 38907S |
| GhostDye 780 | 780 | 1:1000 | Cytek | - | 13-0865-T500 |
| IR Viability | IR | 1:1000 | Fisher Scientific | - | L34982 |
| CD8a | FITC | 1:100 | Biolegend | RPA-T8 | 301006 |
| CD45RA | APC | 1:100 | Biolegend | HI100 | 304112 |
| CD62L | BV421 | 1:100 | Biolegend | DREG-56 | 304827 |
| CD25 | APC | 1:100 | Biolegend | BC96 | 302610 |
| CD69 | BV605 | 1:100 | Biolegend | FN50 | 310938 |
| TIM3 | BV785 | 1:100 | Biolegend | F38-2E2 | 345032 |
| LAG3 | FITC | 1:100 | Biolegend | 11C3C65 | 269308 |

**Supplementary Table 1.** Antibodies used for flow cytometry readout.
